# Fabaceae and Cerrado savanna: Two descriptors of Brazilian extrafloral nectary plants

**DOI:** 10.1101/2024.04.05.588328

**Authors:** Vanessa Dayane da Costa Barbosa, Alexandra Bächtold, Kleber Del-Claro, Estevao Alves da Silva

## Abstract

Extrafloral nectaries (EFNs) have been described in almost 4000 plant species, but there are several gaps in our knowledge of their occurrence and distribution. Here, we investigated the geographical distribution and richness of EFN–plants in Brazilian biomes (Cerrado, Atlantic Forest, Caatinga, Amazon, Pantanal, and Pampa). Data were extracted from 170 papers, and our analysis included only EFN–plants that interacted with ants. A total of 224 EFN–plant species in 115 genera and 48 families were registered in five biomes: Cerrado, Atlantic Forest, Caatinga, Pampa, and the Amazon. The Cerrado was evaluated in 64% of all publications, had the highest richness (90 species) and the most exclusive flora compared to the other biomes. In addition, the most studied species belonged to the Cerrado flora (e.g., *Caryocar brasiliense*). Fabaceae was the most speciose family, with 76 species, being dominant in all biomes and greatly surpassing other families. Only Fabaceae and Euphorbiaceae were found in all biomes, and in general, each biome had its own flora, as only 18 (of 224) plants were found in more than one biome. In a network analysis, *Qualea grandilflora* and *Plathymenia reticulata* were relatively more important than other species, as they connected biomes and increased the cohesion of the community. Our study shows that our understanding of EFN–plants is limited because the Fabaceae and Cerrado were overrepresented. A detailed record of species occurrence and distribution could be a valuable tool for studying the biodiversity of EFN–plants and their potential mutualistic interactions.

## Introduction

Extrafloral nectaries (EFNs) have long served as study models for mutualistic interactions (Bentley 1977). These structures are located mainly on the leaves and release a carbohydrate-rich solution with a few amino acids (Shenoy et al. 2012). Several studies highlight that the evolution of EFNs is especially related to the attraction of ants, which are rewarded with sugars and in turn, act as plants’ bodyguards (Del-Claro et al. 2016, Monique et al. 2022).

EFNs have been described in almost 4000 plant species (Weber & Keeler 2013) and are widely distributed throughout the world, especially in the neotropics (Díaz-Castelazo et al. 2013, Miranda et al. 2022). In fact, the first investigations of EFN–plants and mutualistic interactions with ants have been conducted in the neotropics, particularly in Costa Rica and Panama (Janzen 1969, Bentley 1976). Since then, EFN–plants have been studied in many parts of the globe, but several others remain largely unexplored (reviewed by Juárez-Juárez et al. 2023), thus creating gaps in our knowledge of species occurrence and distribution.

The knowledge of EFN–plants (and their potential interactions with ants) may become biased when research is geographically limited because a few areas are not broadly representative of a biome, region or country (Oliveira et al. 2017). This might create disproportions in the understanding of the processes underlying patterns of distribution and even influence how we prioritize new areas to be explored. A detailed cataloguing of species occurrence and distribution may be an important tool for knowledge of the biodiversity of EFN–plants and potential mutualistic interactions (Aguirre et al. 2013, Nogueira et al. 2015). In addition, it may also indicate gaps in the occurrence of these plants and permit a concentration of efforts to sample unexplored areas (Juárez-Juárez et al. 2023).

When studying the occurrence of species on a large scale, scientists usually use a biome approach (Oliveira et al. 2016), which can be reasonable when dealing with plants (Woodward et al. 2004, Oliveira et al. 2016). Biomes are characterized by their geography, geology and environment, which together create the necessary conditions for the development and persistence of the associated flora (Roesch 2009). Although some overlaps between species may exist along biomes’ boundaries (Pott et al. 2011), most species find their ideal environment in specific biomes, where they have evolved functional characteristics that permit their adaptation to soil, climate conditions and ecological interactions (Pennington et al. 2004, Reu et al. 2011). For instance, plants from the Brazilian savanna are mostly shrubs and short trees that have adapted to fire, while in the Amazon plants may reach tens of meters in height and are not fire tolerant (Hoffmann et al. 2012, Balch et al. 2015).

Recently, there has been a claim about the importance of knowing the geographical distribution of plant–ant mutualisms, as it can synthesize scattered information and reveal unknown spatial patterns that allow further comparisons and questions still needing research (Juárez-Juárez et al. 2023). In the present study, we aimed to map the geographical distribution of EFN–plants that have served as study subjects for plant–ant investigations in Brazilian biomes. Being the largest country in the neotropics, not only does Brazil host many biomes, but it also sustains most studies of EFN–plants (Juárez-Juárez et al. 2023), permitting us to make comparisons and find patterns within plant distributions. In addition, Brazil has two hotspots, the Cerrado and Atlantic Forest, and both have had a consistent tradition of serving as fieldwork for plant–ant studies (Oliveira et al. 1987, Morellato & Oliveira 1994, Calixto et al. 2021).

By using data from published literature, we compiled information about where EFN– plants have been studied in Brazil. Since we were interested in plants that have the potential to establish a mutualistic relationship with ants, we concentrated on species that were reportedly visited by ants. We specifically searched for the geolocation of the fieldwork, and the identity of plant species and families in each biome. With this data, we *(i)* depicted the geographical distribution of where EFN–plants were investigated on a biome scale; *(ii)* examined the most studied biomes, plant species and families; *(iii)* investigated whether plant richness in each biome was related to the number of publications; *(iv)* analyzed the similarity and exclusivity of plants among biomes; and *(v)* implemented a network analysis approach to evaluate the most important EFN–plants (species and families) in each biome and the importance of each biome in supporting EFN–plants.

## Methods

### Brazilian biomes

Brazil is located in South America and sustains six terrestrial biomes, namely the Amazon, Cerrado (Brazilian savanna), Atlantic Forest, Caatinga, Pampa, and Pantanal (**Figure 1**). Two of them, the Cerrado and Atlantic Forest, are regarded as hotsposts (Myers et al. 2000). These six biomes present over 32,000 species of angiosperms in 229 families (Zappi et al. 2015). Some brief characteristics of each biome are provided in **Supplementary File 1.**

**Figure 1.**
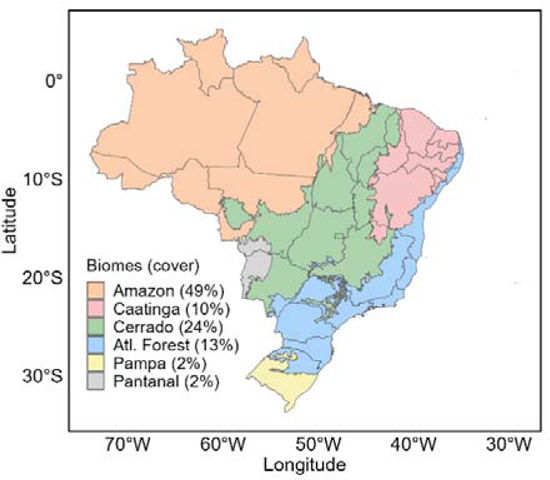
Brazilian map showing the biomes and their respective land cover (see the names of states in **Supplementary File 2**).

### Data collection

The search for literature on EFN–plants was performed using *Google Scholar* as the main search tool, as this platform provides outputs from different journals and often permits papers to be downloaded for free. *Google Scholar* may also forward the search to third-party websites, where the papers can be downloaded. In a preliminary search comparison, other databases, such as *Scielo* and *Web of Science*, provided the same results as *Google Scholar*; thus, only the latter was used.

The search for papers was conducted in four steps. First, we used the words “extrafloral nectar*” and “Brazil” (with Portuguese variations, such as nectários extraflorais and Brasil). After a collection of several papers, we conducted a second search using the entries above and the names of each biome. The third step consisted of conducting a search using the term “extrafloral nectar*” only. To conclude, in the fourth step, we scrutinized the references’ list of papers to gather additional publications that might not appear in the internet search. We restricted the data to papers published until the end of 2022. Papers were considered published as soon as they appeared in the journal’s websites (e.g., in press, early access, early view, online first) (Alves-Silva et al. 2016).

The literature search commenced in mid-2021 and lasted until the last quarter of 2023, when we came across new studies published in 2022 that were not available in previous searches (Barbosa 2023). Thus, we considered our data to cover as many published papers as possible. Papers that were not available for download were requested from the authors.

### Data criteria

Some conditions were established for a paper to be included in our database: ***(i)*** papers with geographical information on the fieldwork, as it permitted us to determine the biome where the plants were located. Studies conducted in botanical gardens, greenhouses, or laboratories were not considered, as plants might be out of their natural range of occurrence. ***(ii)*** Papers published in journals with editorial boards and a history of peer review. ***(iii)*** Plants that were observed to interact with ants. ***(iv)*** Papers with plant species identification. Since we were interested in relating plants with different biomes, as well as classifying plants as important for the community, we inevitably required the identification of plants to the species level.

### Considerations of the criteria

Our interest in this study was not to provide a checklist of EFN–plants in Brazil, but rather to show the plants in different biomes that were indeed visited by ants and have served (and can serve) as study subjects for plant–ant investigations. The choice to use only EFN– plants visited by ants permitted us to concentrate on plants that might establish potential mutualism interactions. We understand that the simple presence of ants on plants is not evidence of mutualism (Nogueira et al. 2012), but at least ants visited the EFNs of identified/known plants. Thus, scientists have a starting point to investigate potential mutualisms.

By examining only plant species with full identification, we might have excluded a portion of the EFN–flora. However, the lack of species identification might inhibit our analyses, as we were interested in plant distributions in relation to biomes. A few studies provided the information of genera only, and this information was not used. Some genera can have continental distribution, e.g., *Inga* (Kersch & Fonseca 2005, Domingos & Alves Silva 2023), thus not contributing to our understanding in a biome approach. Another problem in dealing with genera is that the outcomes of mutualism may present variations. For instance, five species of *Banisteriopsis* (Malpighiaceae) have been studied regarding interactions with ants, and the results are species specific (Alves-Silva & Del-Claro 2016, Mendes-Silva et al. 2024). In addition, one issue with *Banisteriopsis* is that it is one of the richest genera in the Cerrado and may be misidentified with other genera, some of which do not have EFNs (Araújo et al. 2020). Additionally, in recent years, many plant species have been incorporated into other genera (e.g., *Distictella elongata* (Vahl) Urb is now *Amphilophium elongatum* (Vahl) L.G. Lohmann) and even in other families. Thus, a conservative approach that considers only plant species seems more appropriate. Moreover, we performed network analyses, and most authors use plant species in the analyses (Fagundes et al. 2017, Silva et al. 2020).

We focused only on EFN–plants; the ants were not considered because of taxonomical problems inherent to ant identification, either in the field or in the laboratory (Souza et al. 2024). Consequently, several studies did not provide the identification of ants to species level that were visiting the EFN–plants. Another issue in dealing with ants is that the authors may provide a list of EFN–plants but do not link the ants that were visiting each plant species. Forthcoming approaches should be attempted before considering ants in a similar study.

### Biomes

Defining and classifying biomes can be challenging, but recent advances have improved the concept (Pennington et al. 2004, Faber-Langendoen et al. 2020, Navarro et al. 2023). Biomes have both a global (from desert to tundra) and a regional classification based mostly on rainfall and temperature (Myers et al. 2000, Woodward et al. 2004). For the sake of clarity, we considered the six terrestrial biomes of Brazil recognized by the Ministry of Environment (Cerrado, Amazon, Caatinga, Atlantic Forest, Pantanal and Pampa) (Roesch et al. 2009, Simon et al. 2009, Silva et al. 2017). In addition, most of the literature dealing with EFN–plants describes the occurrence of plants based on biomes and even uses the characteristics of these regions to explain ecological factors and outcomes in plant–ant interactions (Oliveira & Leitao-Filho 1987, Leal et al. 2015, Oliveira et al. 2021, Miranda et al. 2022).

Information on biomes (i.e., where the study was carried out) was extracted from the papers, as the authors mentioned either the biome or the characteristics of the area where the fieldwork was conducted (e.g., terra firme forests, rainforest, semi-arid, semideciduous forest). In transition zones between biomes, the authors specify the dominant vegetation where the study was conducted (e.g., rupestrian fields, which is a Cerrado phytophysiognomy), and the biome was labeled accordingly (Monique et al. 2022). In cases where the biome was not mentioned, and there was no similar fieldwork to compare with, we checked the IBGE (Instituto Brasileiro de Geografia e Estatística) website (https://cidades.ibge.gov.br), as it contains data on the biomes of each Brazilian municipality. The authors of the papers were also consulted.

### Extrafloral nectary definition

Extrafloral nectaries are regarded as organs in vegetative structures that produce water, carbohydrates and other substances, and are not related to pollination (Weber & Keeler 2013, Marazzi et al. 2019) (Figure 2). Nonetheless, some scientists also considered extranuptial and pericarpal nectaries (nectaries present in reproductive structures not related to pollination) to be EFNs (Oliveira-Filho et al. 2022). This occurs because, similar to leaf EFNs, these nectaries are visited by ants that may act as antiherbivore defense (Oliveira-Filho et al. 2022). In the present study, the extranuptial and pericarpal nectaries were also considered EFNs, and publications on this subject were included in our database.

**Figure 2.**
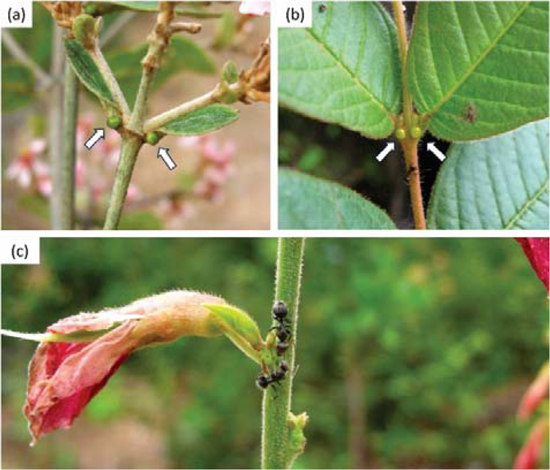
Extrafloral nectaries in (a) *Heteropterys pteropetala* (Malpighiaceae) and (b) *Qualea grandiflora* (Vochysiaceae); (c) ants visiting the extrafloral nectaries of *Bionia coriacea* (Fabaceae).

### Data organization

We retrieved data from 170 papers published in 70 journals. We extracted the following information: journal name, authorship, year of publication, plant species, study site (park, reserve, university campus), city of fieldwork, state, biome and geolocation (latitude and longitude) of fieldwork. Given the existence of several fieldwork locations, including parks, reserves, and urban areas, we grouped them all into cities for the sake of clarity and to ease the points of occurrence in maps. When the fieldwork was conducted in parks or reserves, we used *Google Earth* as a tool to examine the nearest cities, as well as the internet, to determine to which city a park or reserve was associated. Plant taxonomical identification was checked on the websites of Flora do Brasil (http://floradobrasil.jbrj.gov.br/) and Sistema de Informação sobre a Biodiversidade Brasileira (https://sibbr.gov.br/).

### Analyses

Descriptive data are shown as means and standard deviations. The geographical distribution of fieldwork was plotted on a map with Brazil boundaries overlaid with all six biomes (*objective i*). Figures were generated in R statistical software using the packages ggplot2, geobr, and sf (Wickham 2016, Pereira & Gonçalves 2022, Pebesma & Bivand 2023). Chi-squared tests were performed to compare the frequency of studies in each biome, the most studied plant species and families, and the richness of plants and families per biome (*objective ii*). Since our data contained over 200 plant species, we decided to include in this analysis only those species that were sampled more than five times.

The relationship between plant species (dependent variable), plant families, biomes and the number of publications (factors) was investigated with a general linear model test and Poisson distribution (*objective iii*). In this test, the interaction effect was not considered, as it was not significant and overdispersed the model.

The relation between plant species (dependent variable), plant families and biomes (factors) was investigated with a general linear model test, with Poisson distribution (*objective iii*). In this test, the interaction effect was not run as it was not significant and over dispersed the model. The Jaccard dissimilarity index was used to compare the plant richness among biomes (*objective iv*), and the number of exclusive species and families in biomes was compared with chi-squared tests.

To compare the importance of each biome in sustaining plant species and families, we built ecological networks (*objective v)*. Networks were composed of pairwise interactions between plants (species and families) and biomes (both regarded as nodes) built under a matrix of presence and absence, in which the connection between both parties indicates a link (Trøjelsgaard & Olesen 2016, Guimarães 2020). With this organization of data, we identified the biomes that had the most records of EFN–plants and which nodes (plant species, plant families and biomes) were the most important for network cohesion.

To measure this in numbers, we calculated the betweenness centrality index, as it provides an appropriate estimation of the most important nodes in a network, not only by giving weight to plants and biomes with most interactions, but also by considering plants that connect different biomes and, if removed, cause the rupture of the network (González et al. 2010, Sazima et al. 2010, Farine & Whitehead 2015, Trøjelsgaard & Olesen 2016). Betweenness centrality is unitless, but its interpretation is simple, as nodes with higher values are regarded as the most important for the community. For instance, biomes with the highest values are important both because they have many associated plants and because they share some of the flora with other biomes. For plants, a high betweenness value indicates that they are important connectors among biomes and that their absence could create modules, i.e., biomes without connections with others.

Two networks showing the betweenness centrality values were created, the first for all plant species and biomes, and the second showing all plant families and biomes. In these networks, we determined the relative importance of each biome, the plant species that occurred in more than one biome, and the most important species and families in the EFN plant community.

Another set of four networks was generated using data from the most speciose families, namely Fabaceae, Bignoniaceae, Euphorbiaceae, and Malpighiaceae. Together, these families accounted for 61% of species (*n* = 136) and were present in 58% (*n* = 81) of publications. In these networks, we calculated the degree centrality index, which was simply a depiction of how many species within each family were present in each biome. The betweenness centrality was not calculated here, as the previous networks already had the purpose of showing which families were relatively more important to the community. In these networks with Fabaceae, Bignoniaceae, Euphorbiaceae, and Malpighiaceae, we intended to show the biomes that sustained more species of these families. Networks were generated with package *igraph* in R (Csardi & Nepusz 2006).

Network analyses can be accompanied by a multitude of metrics, but we decided to simplify and calculate just a few, both to keep the study as clean as possible and because many of the network metrics were related, thus generating redundancies (Sazima et al. 2010, Delmas et al. 2019, Souza et al. 2024). Data from the Pampa biome were not used in the networks because of the low sample size (*n* = 2 plant species and both from a single family).

## Results

### Points of occurrence

Plants were sampled in five biomes (Cerrado, Atlantic Forest, Caatinga, Amazon, and Pampa; Pantanal was not represented), 18 federated units (including the Federal District, the capital of Brazil), and 62 cities (Figure 3). The distribution of fieldwork revealed that most studies were conducted in the Cerrado, Atlantic Forest (above 23° South latitude) and Caatinga biomes. This study-belt accounted for 83% (*n* = 51) of all points of occurrence.

**Figure 3.**
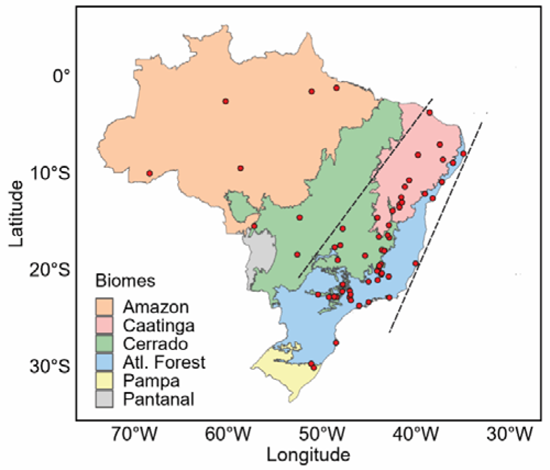
Geographical distribution (points of occurrence) of where extrafloral nectary plants were sampled. A given point of occurrence may indicate one or more publications in the same city. Dashed lines delimit a concentration of sampling areas.

The Cerrado biome was evaluated in 64% (*n* = 109) of all publications, followed by the Amazon (15%, *n* = 25), Atlantic Forest (11%, *n* = 19), Caatinga (9%, *n* = 15), and Pampa (1%, *n* = 2) (□^2^ = 162.62, *df* = 3, *P* < 0.0001). Only six papers (3.5% of all data) investigated EFN-plants in more than one city, but always in the same biome.

### Plant richness

Plant richness accounted for 224 species in 115 genera and 48 families (**Supplementary File 3)**. The richness of the plants investigated in each paper ranged from 1 to 28 species (3.27 ± 5.62 plants per paper), but most papers (*n* = 124; 73%) investigated only one plant species. The number of plant families per paper ranged from 1 to 13 (1.91 ± 2.37), with most studies (*n* = 136; 80%) investigating one family.

Fabaceae was present in 48 papers (38% of total) (□^2^ = 697.77, *df* = 37, *P* < 0.0001) and was also the richest family with 76 species (□^2^ = 1137.2, *df* = 39, *P* < 0.0001). This was more than two-fold the richness of the second most studied family, the Bignoniaceae, with 32 plant species (Figure 4**, Supplementary File 4)**. Fabaceae was the richest family in all biomes (Figure 5).

**Figure 4.**
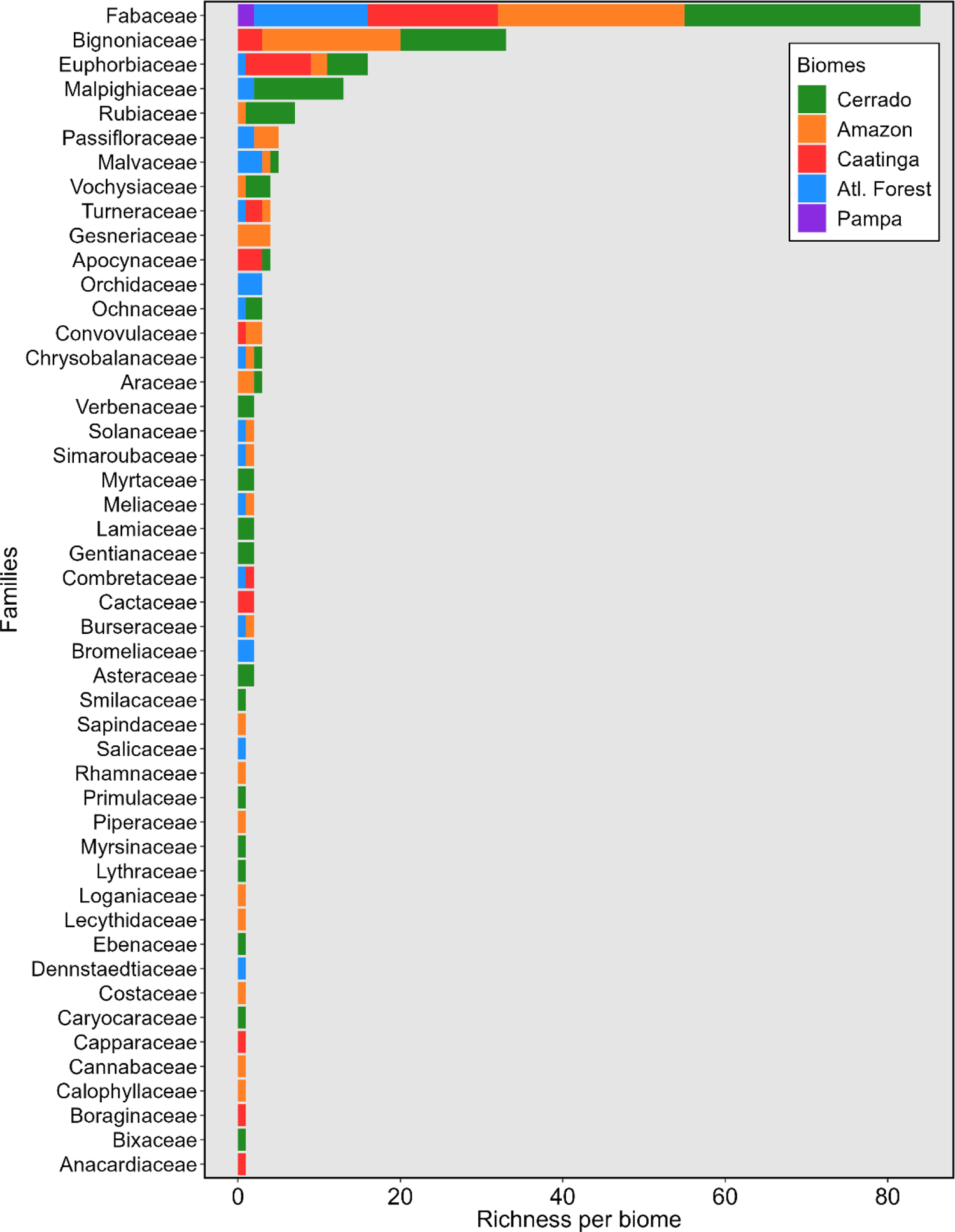
Richness of extrafloral nectary plants according to families and biomes.

**Figure 5.**
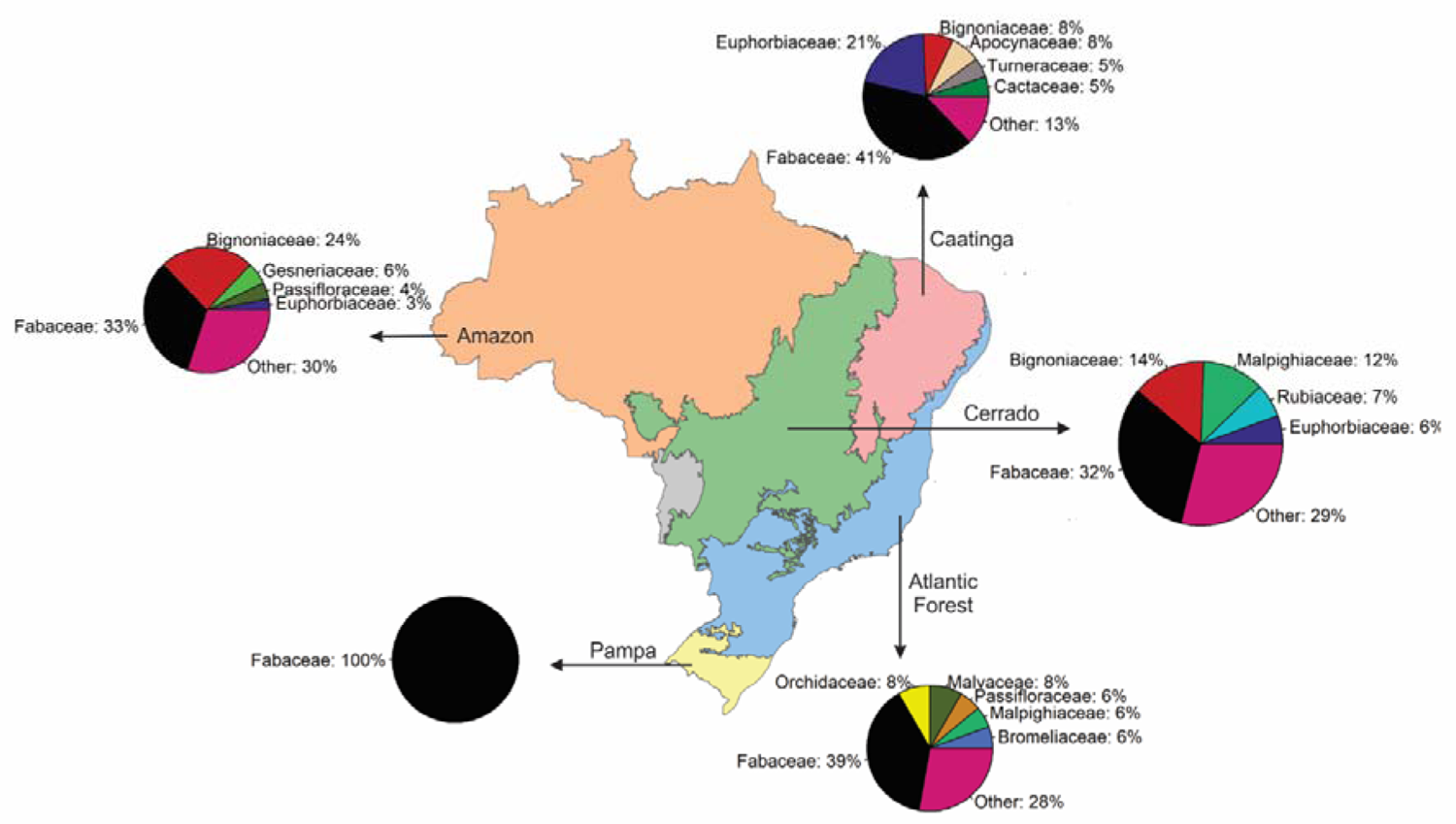
The most representative plant families according to biomes. Note that Fabaceae had the highest richness in all biomes.

The Cerrado was overrepresented in terms of plant species (□^2^ = 95.91, *df* = 3, *P* < 0.0001) (Figure 6a**, Supplementary File 5**). However, the number of families was higher in the Amazon (*n* = 24 versus 23 in the Cerrado) (Figure 6b). Plant richness was influenced by the biome (Cerrado had the most species) and family (Fabaceae was far more speciose than the other families). The number of papers did not affect the richness of EFN–plants (**Table 1**).

**Figure 6.**
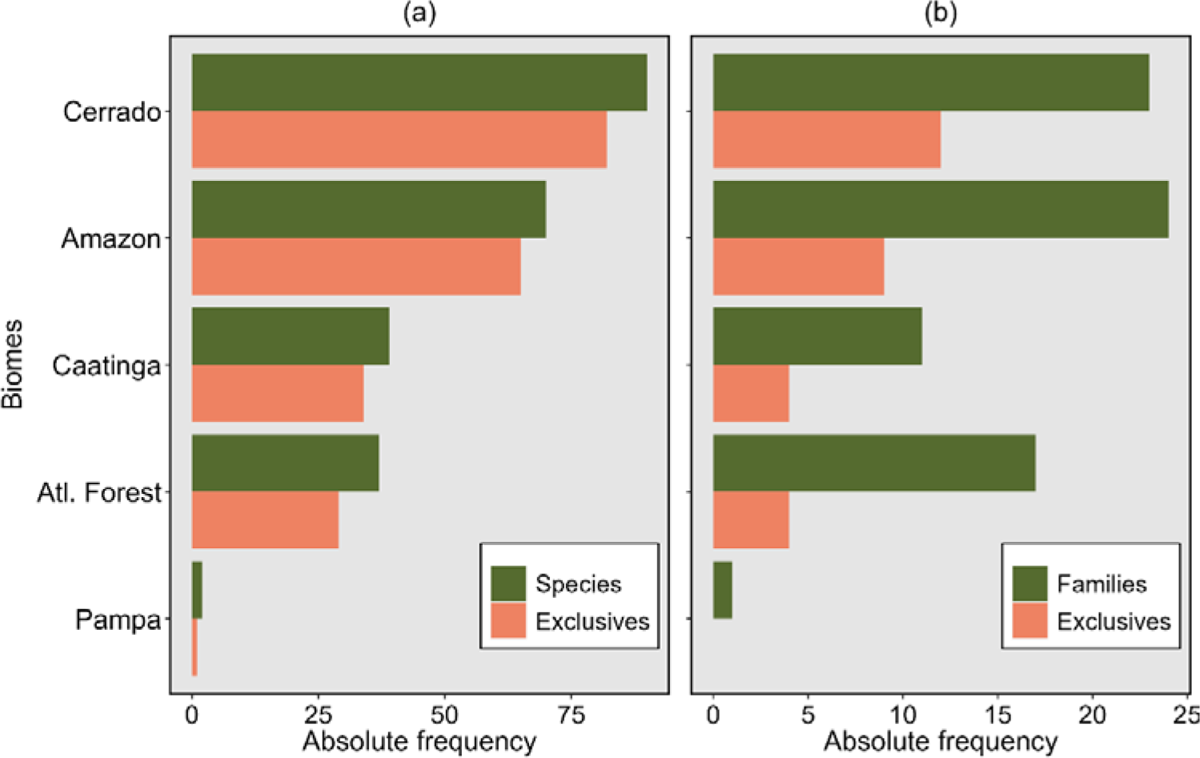
**(a)** Total number of species and exclusive species of extrafloral nectary plants sampled in each biome; **(b)** the same rationale, but for families. The Pampa had no exclusive families as only Fabaceae was sampled in this biome, and this family occurred in all the other biomes.

**Table 1.**
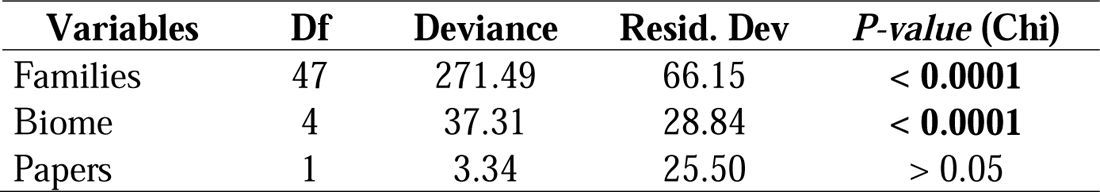
Factors influencing the richness of extrafloral nectary plants. Interaction effects were not incorporated in the model as they were no significant.

### Exclusive species

The Cerrado had the most exclusive pool of species and families (Figure 6a**, b; Supplementary File 6**). However, the results were not significant (□^2^ tests, *P* > 0.05 in both cases), as exclusivity depended on the actual richness, i.e., more species led to a higher likelihood of retaining more exclusive species (**Supplementary File 6)**.

### Most studied plants

Most plants were investigated only once (53%) or twice (23%), and only a small group of 20 plants (out of 224 species) was present in more than five papers. The most studied EFN plant was *Caryocar brasiliense* Cambess (Caryocaraceae) (□^2^ = 70.36, *df* = 19, *P* < 0.0001), followed by several other plants from the Cerrado biome. In fact, of the 20 top-studied plant species, only two were not present in the Cerrado (Figure 7).

**Figure 7.**
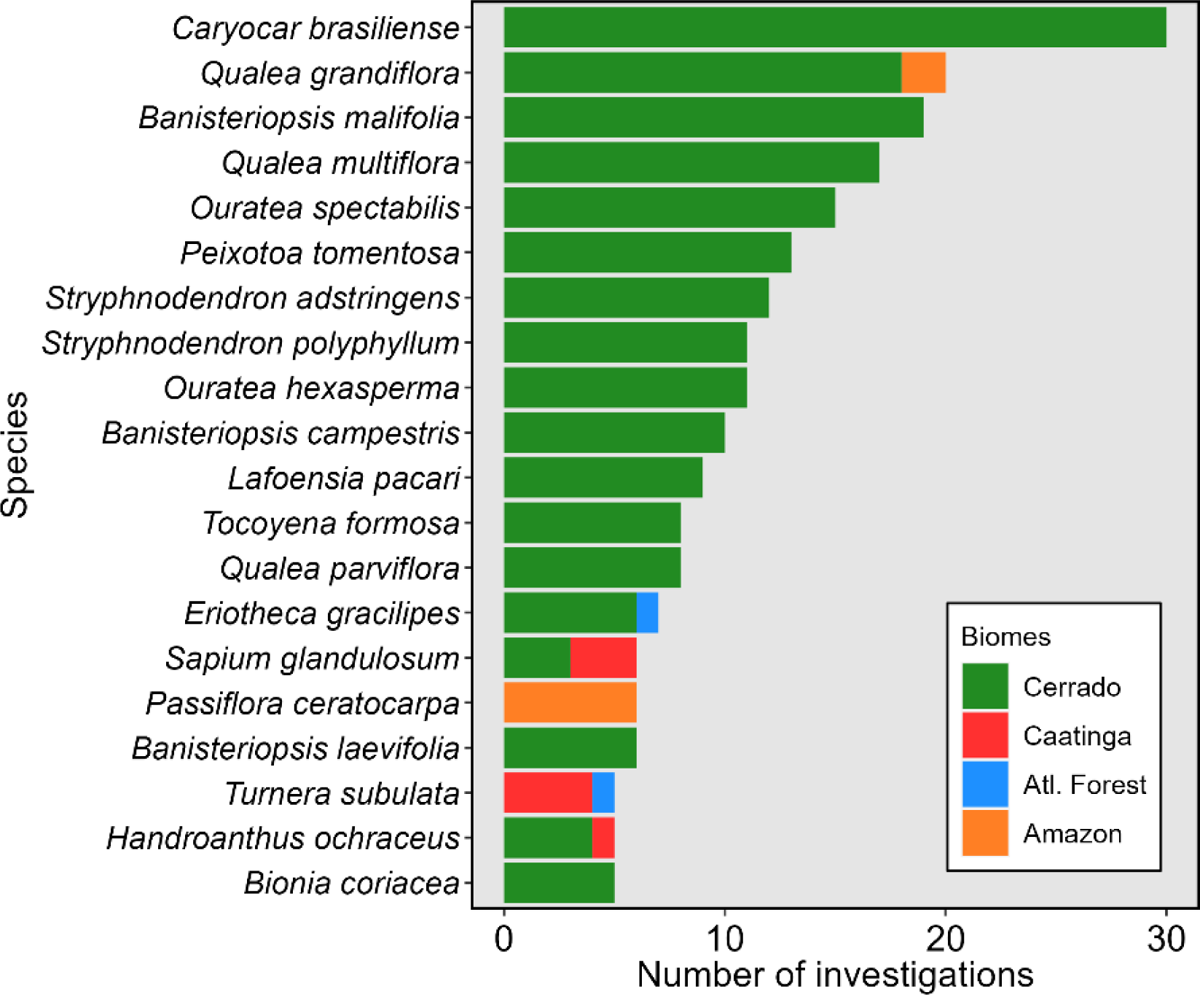
Extrafloral nectary plant species according to biomes. The figure shows plants that were investigated ≥ 5 times to allow comparisons. The Pampa biome was not represented as the sample size was low.

### Flora similarity

Only 12 plant species (out of 224) and 19 families (out of 48) occurred in more than one biome. No plant species was registered in all biomes. Regarding families, only Fabaceae was found in all five biomes (all the two plant species sampled in Pampa belonged to Fabaceae), followed by Euphorbiaceae (Figure 4). In general, the dissimilarity of flora among biomes was high (**Table 2**).

**Table 2.**
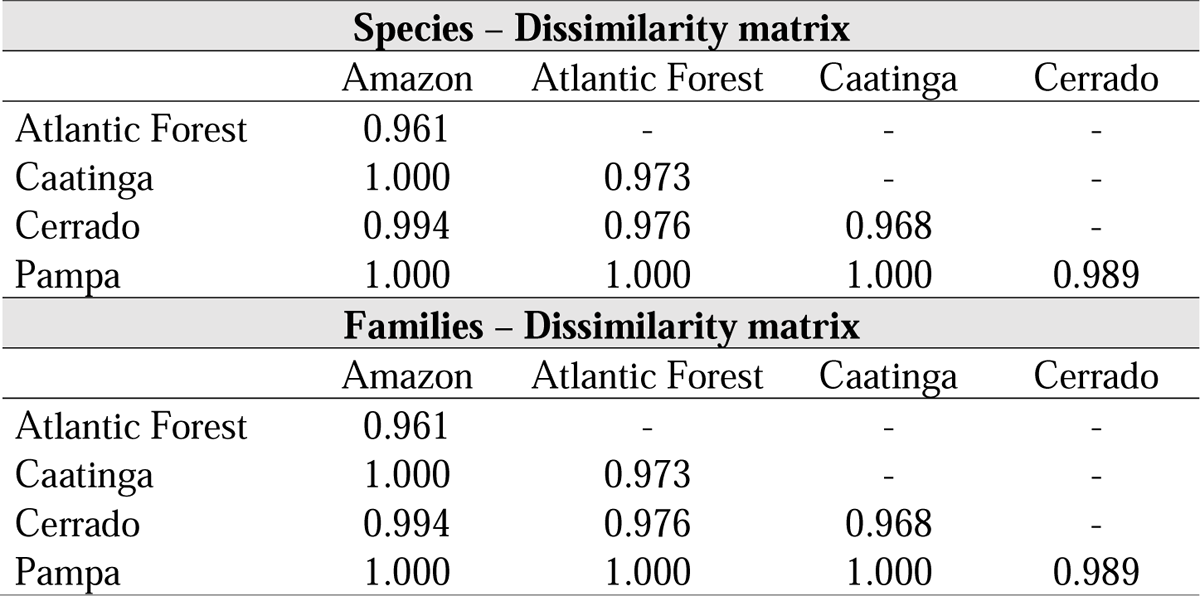
Jaccard dissimilarity matrix with data of plant richness and number of families in each biome where extrafloral nectary plants were sampled.

| **Species – Dissimilarity matrix** |  |  |  |  |
| --- | --- | --- | --- | --- |
|  | Amazon | Atlantic Forest | Caatinga | Cerrado |
| Atlantic Forest | 0.961 | - | - | - |
| Caatinga | 1.000 | 0.973 | - | - |
| Cerrado | 0.994 | 0.976 | 0.968 | - |
| Pampa | 1.000 | 1.000 | 1.000 | 0.989 |
| **Families – Dissimilarity matrix** |  |  |  |  |
|  | Amazon | Atlantic Forest | Caatinga | Cerrado |
| Atlantic Forest | 0.961 | - | - | - |
| Caatinga | 1.000 | 0.973 | - | - |
| Cerrado | 0.994 | 0.976 | 0.968 | - |
| Pampa | 1.000 | 1.000 | 1.000 | 0.989 |

### Network analyses

The Cerrado was the most important biome in terms of EFN-bearing plants, not only because it sustained many species, but also because it shared many species with all the other biomes (Figure 8a). Among the plant species, a higher betweenness value was noted for *Qualea grandiflora* Mart. (Vochysiaceae), the sole plant linking the Amazon and Cerrado biomes. The only plant sampled in three biomes was the *Plathymenia reticulata* Benth. (Fabaceae), and consequently, it had a high betweenness value (Figure 8a). Of the 12 species that occurred in more than one biome, six were Fabaceae.

**Figure 8.**
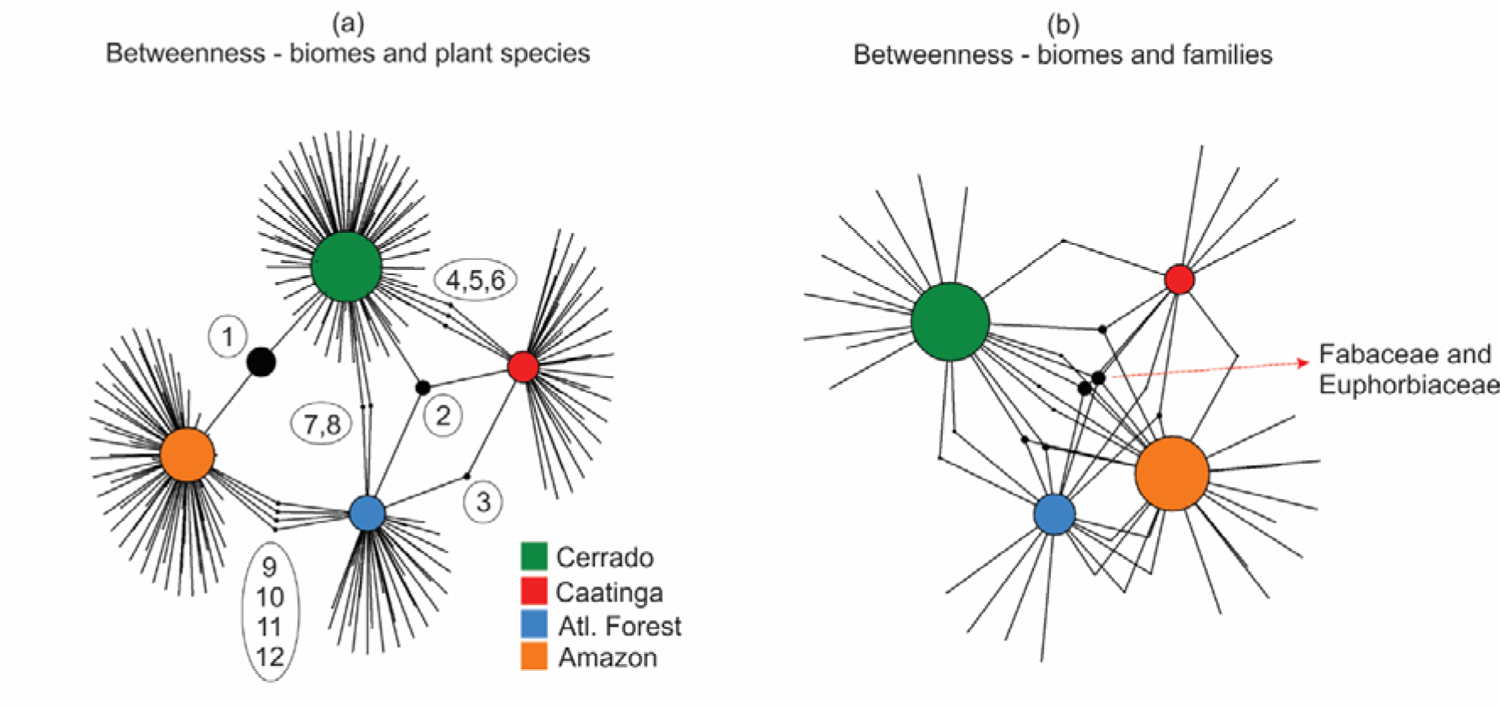
Network indicating the most important **(a)** plant species and **(b)** families for the cohesion of the extrafloral plant community in all biomes. Larger circles indicate the most important nodes for the network. The names in **(a)** are 1 - *Qualea grandiflora,* 2 - *Plathymenia reticulata*, 3 - *Turnera subulata,* 4 - *Handroanthus ochraceus,* 5 - *Senna velutina*, 6 - *Sapium glandulosum*, 7 - *Copaifera langsdorfi,* 8 - *Eriotheca gracilipes,* 9 - *Stryphnodendron pulcherrimum,* 10 - *Inga capitata,* 11 - *Inga edulis,* 12 - *Simarouba amara*.

The most important families were Fabaceae and Euphorbiaceae, as they were present in all biomes (remind that data of Pampa were not used because of the low sample size) (Figure 8b). The Amazon and the Cerrado had the highest betweenness values, evidence of their importance in sustaining many families and sharing these families with other biomes.

The richest families (Fabaceae, Bignoniaceae, Euphorbiaceae, and Malpighiaceae) showed a distinct distribution among the biomes. Fabaceae was distributed in all biomes; Bignoniaceae was the richest in the Amazon, and Euphorbiaceae had most species in the Caatinga. Malpighiaceae was almost exclusive to the Cerrado (Figure 9). Fabaceae shared six species with other biomes, while Bignoniaceae and Euphorbiaceae shared one (Figure 9).

**Figure 9.**
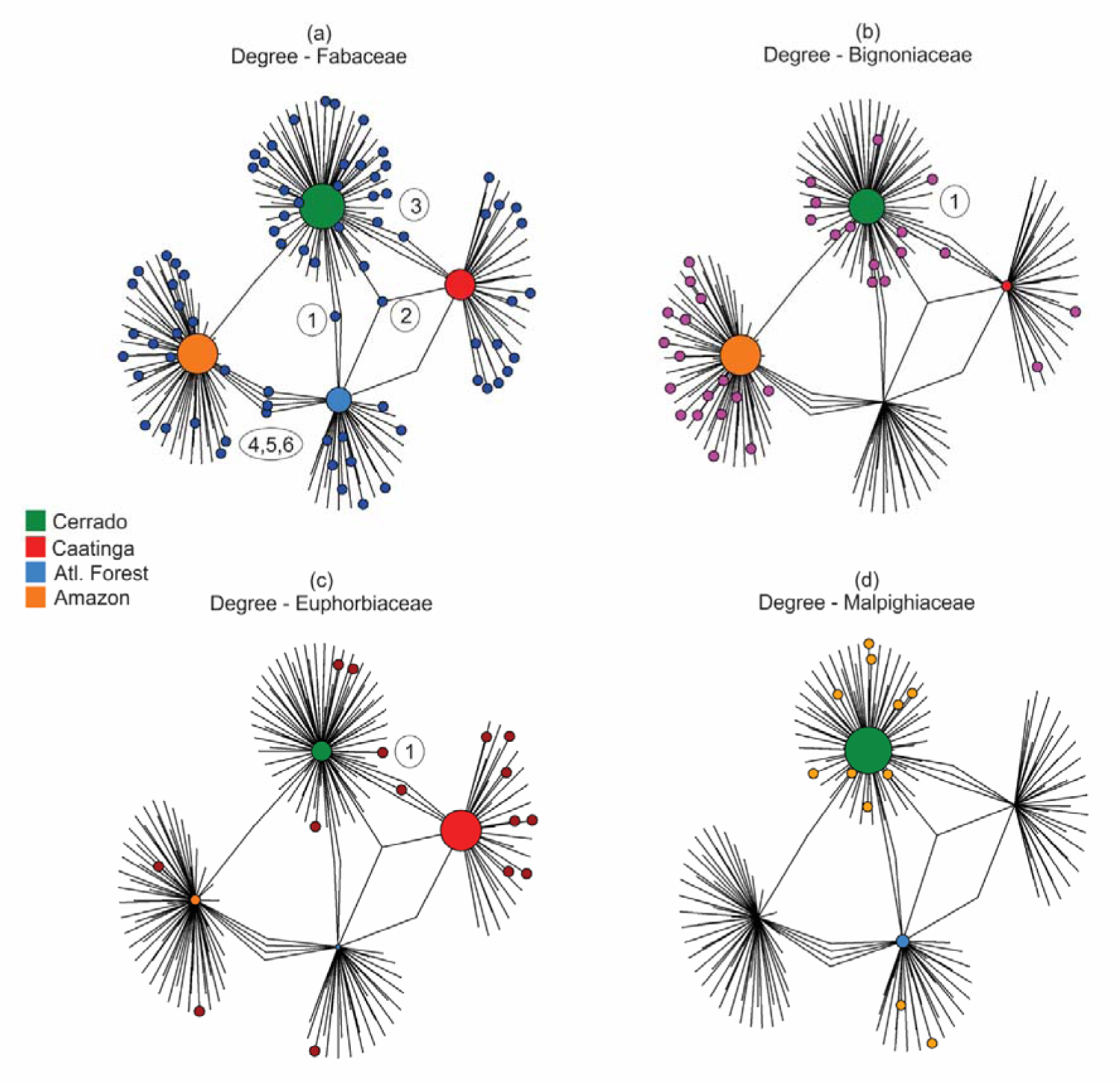
Degree distribution of the extrafloral nectary plants in four families according to biomes: **(a)** Fabaceae; **(b)** Bignoniaceae; **(c)** Euphorbiaceae; **(d)** Malpighiaceae. These were the richest families of extrafloral nectary plants. The terminal-colored dots indicate the richness in each family, while the lines without colored dot indicate species in other families. The size of the biomes’ circles indicates the relative richness of plants the biome is associated with. Species of Fabaceae that occurred in more than one biome **(a)** 1 – *Copaifera langsdorffii*, 2 – *Plathymenia reticulata*, 3 – *Senna velutina*, 4 - *Inga capitata*, 5 - *Inga edulis*, 6 – *Stryphnodendron pulcherrimum*. Species of Bignoniaceae **(b)** 1– *Handroanthus ochraceus*. Species of Euphorbiaceae **(c)** 1 – *Sapium glandulosum*.

## Discussion

### Publications, plants, and biomes

The Cerrado biome accounted for over 60% of all publications on EFN–plants, had the largest plant richness, was second place in terms of families, and had the most exclusive pool of species and families. The most frequently studied plant species was the Cerrado native *C*. *brasiliense*, which was present in 30 publications. Conversely, most of the other 223 species were studied only once or twice. In addition to *C*. *brasiliense*, most of the other most frequent species were from the Cerrado. At first sight, this might be a problem and show a bias in our knowledge of plant–ant interactions. In fact, it is surprising that among a pool of several other co-occurring EFN–plant species in the field, scientists tend to choose the *C*. *brasiliense* for investigation. Areas where *C*. *brasiliense* was sampled sustains several other EFN–plant species (Sendoya et al. 2009, Lange et al. 2013, Moura et al. 2022). The other top-studied plants such as *Qualea*, *Banisteriopsis* and *Ouratea* also occur in a community with several other EFN–plants.

One reason to explain the predominance of a few species as study subjects is that once a study-system is established scientists tend to explore it in several ways. Since plant–ant interactions are highly conditional over time and space and are influenced by biological factors (ants and herbivores) (Franco & Cogni 2013, Koch et al. 2016, Del-Claro et al. 2016, Cruz Rocha et al. 2019, Domingos & Alves Silva 2023, Souza et al. 2024), studying several aspects of this mutualism in a few species might provide rich data, allowing for comparisons among different approaches, time, locales, and associated arthropod fauna.

In this context, by using *C*. *brasiliense* (and other frequently studied plants) as a study model, scientists have been able to unveil several aspects of the ecology of the plant and its interactions with ants and herbivores (Oliveira 1997, Byk & Del-Claro 2010, Neves et al. 2012, Camarota et al. 2015, Moura & Del-Claro 2023). For many plant species, the outcomes of the interaction with ants are unknown. Thus, the investigation of some plant species is justified by the amount of information that can be extracted from the ecological system.

### Important nodes (plants and biomes)

Based on network analyses, both the Cerrado and Fabaceae can be considered important for the EFN–plant community. Classifying a node (species, families and biomes in our study) as important or key may be a difficult and arbitrary task because ecological interactions are complex. Therefore, scientists often use analytical techniques to emphasize species that stand out in a system (Costa et al. 2016, Cagua et al. 2019). The centrality indexes used in this study are based on both the direct and indirect connections among plants and biomes, and they also give weight to plants that bonded to different biomes, and to the biomes that shared plant species with other biomes (Farine & Whitehead 2015, Souza et al. 2024). For instance, the Caatinga had a higher degree (more plants) than the Atlantic Forest, but the latter had higher betweenness because its plants connected all biomes. In terms of degree, the Cerrado stood out as the biome with the most pairwise interactions (highest plant richness) in relation to the other biomes. In addition, since the Cerrado shared many species with the other biomes, it also had high betweenness.

Fabaceae was the most important family in the network because it had a high richness, was present in all biomes, and contained many species that connected biomes. Euphorbiaceae was also relatively important in the network, but it was less rich than Fabaceae. *Caryocar brasiliense* can be considered one of the most important species in the network, as it was the most studied species; nonetheless, it is restricted to the Cerrado biome. Conversely, less studied species, such as *Q*. *grandiflora* and *P*. *reticulata*, had higher betweenness because they connected different biomes. In this sense, although *C*. *brasiliense* is important to the understanding of plant–ant interactions in the Cerrado, *Q*. *grandiflora* and *P*. *reticulata* may provide information from other biomes. In addition, should *Q*. *grandiflora* be removed from the network, the Cerrado and the Amazon would lose their connection (Farine & Whitehead 2015). Species with high values of betweenness tend to participate in several trophic chains and affect species abundance and occurrence (Jordán et al. 2007); thus, the plants common among biomes can be a link between the ant fauna (and other associated arthropods) in different biomes (González et al. 2010).

### Exclusive flora

Only a few plant species occurred in more than one biome, and this result is not entirely unexpected. Plants have evolved several adaptive characteristics that permit them to persist in specific conditions of soil, water, climate, and animal partners (Woodward et al. 2004, Hoffmann et al. 2012). Nonetheless, the number of exclusive families was surprising. In the Cerrado, for instance, about half of the families of EFN–plants did not occur in any other biome. These results, however, must be seen with careful consideration. Some plants labeled in our study as exclusive of some biomes may actually be present in other biomes. This occurred because we focused on plants that were reportedly visited by ants. For instance, we found ants visiting *Tocoyena formosa* (Cham. & Schltdl.) K. Schum (Rubiaceae) in the Cerrado, but this plant also occurred in the Atlantic Forest, where interactions with ants have not been reported (Queiroga & Moura 2017, Miranda et al. 2022). By classifying plants and families as exclusive, we can determine to what extent plant–ant interactions have been studied on a large scale.

### Details of the study

EFN-bearing plants are distributed among over 300 families, 90 families, and almost 4000 species (Marazzi et al. 2013, Weber & Keeler 2013, Weber et al. 2015). The numbers in our study may seem noticeably low (48 families, 115 genera, and 224 species) for a country with continental dimensions, such as Brazil. Nonetheless, we investigated only plants that sustained ants because we were interested in plants that might potentially interact mutualistically with ants. Thus, checklists of EFN–plants were left out (Miranda et al. 2022). Should one consider the EFN plant flora of Brazil as a whole, these numbers are supposed to increase considerably.

## Conclusions

One goal of the United Nations Sustainable Development is “*to protect*, *restore*, *and promote the conservation and sustainable use of terrestrial ecosystems*”. The first step is knowledge about where biodiversity occurs. Thus, studies of plant distribution can be a first step to understanding and conserving biodiversity, infer key processes for the maintenance of interactions, and permit assumptions of biogeographical and macroecological hypotheses. Our study provided important information about EFN–plants in all Brazilian territories and revealed a concentration of studies in a region that covers the Cerrado, Atlantic Forest, and Caatinga. This restricted range is responsible for most of what we know about EFN–plant locations. Overall, we may consider the Cerrado biome and the Fabaceae as the variables that represent most of what we know about EFN plants.

## Acknowledgments

We thank Mario Guilherme de Biagi Cava, Daniel de Paiva Silva and Rafael Barbosa Pinto for comments on early versions of this work. We also thank the *Instituto Federal de Educação*, *Ciência e Tecnologia Goiano (IF Goiano)* campus Urutaí for their support, as well as the Programa de Pós-graduação em Conservação de Recursos Naturais do Cerrado (CRENAC, IF Goiano, campus Urutaí).

## References

Aguirre, A., Coates, R., Cumplido-Barragán, G., Campos-Villanueva, A. & Díaz-Castelazo, C. 2013. Morphological characterization of extrafloral nectaries and associated ants in tropical vegetation of Los Tuxtlas, Mexico. Flora: Morphology, Distribution, Functional Ecology of Plants 208:147–156.

Alves-Silva, E. & Del-Claro, K. 2016. On the inability of ants to protect their plant partners and the effect of herbivores on different stages of plant reproduction. Austral Ecology 41:263–272.

Alves-Silva, E., Porto, A. C. F., Firmino, C., Silva, H. V., Becker, I., Resende, L., Borges, L., Pfeffer, L., Silvano, M., Galdiano, M. S., Silvestrini, R. & Moura, R. 2016. Are the impact factor and other variables related to publishing time in ecology journals? Scientometrics 108:1445–1453.

Araújo, J. S., De Almeida, R. F. & Meira, R. M. S. A. 2020. Taxonomic relevance of leaf anatomy in *Banisteriopsis* C.B. Rob. (Malpighiaceae). Acta Botanica Brasilica 34:214–228. scielo.

Balch, J. K., Brando, P. M., Nepstad, D. C., Coe, M. T., Silvério, D., Massad, T. J., Davidson, E. A., Lefebvre, P., Oliveira-Santos, C., Rocha, W., Cury, R. T. S., Parsons, A. & Carvalho, K. S. 2015. The susceptibility of southeastern Amazon forests to fire: insights from a large-scale burn experiment. BioScience 65:893–905. Oxford University Press.

Barbosa, V. D. Da C. 2023. Padrão geográfico de plantas com nectários extraflorais em biomas brasileiros. Instituto Federal Goiano, campus Urutaí. 40 pp.

Bentley, B. L. 1976. Plants bearing extrafloral nectaries and the associated ant community: interhabitat differences in the reduction of herbivore damage. Ecology 57:815–820.

Bentley, B. L. 1977. Extrafloral nectaries and protection by pugnacious bodyguards. Annual Review of Ecology and Systematics 8:407–427.

Byk, J. & Del-Claro, K. 2010. Nectar- and pollen-gathering *Cephalotes* ants provide no protection against herbivory: a new manipulative experiment to test ant protective capabilities. Acta Ethologica 13:33–38. Springer-Verlag.

Cagua, E. F., Wootton, K. L. & Stouffer, D. B. 2019. Keystoneness, centrality, and the structural controllability of ecological networks. Journal of Ecology 107:1779–1790.

Calixto, E. S., Lange, D. & Del-Claro, K. 2021. Net benefits of a mutualism: Influence of the quality of extrafloral nectar on the colony fitness of a mutualistic ant. Biotropica 53:846–856.

Camarota, F., Powell, S., Vasconcelos, H. L., Priest, G. & Marquis, R. J. 2015. Extrafloral nectaries have a limited effect on the structure of arboreal ant communities in a neotropical savanna. Ecology 96:231–240.

Costa, F. V., Mello, M. A. R., Bronstein, J. L., Guerra, T. J., Muylaert, R. L., Leite, A. C. & Neves, F. S. 2016. Few ant species play a central role linking different plant resources in a network in rupestrian grasslands. PLOS ONE 11:e0167161.

Cruz Rocha, M. L., Cristaldo, P. F., Santos Lima, P. S., Dos Santos, A. T., Do Sacramento, J. J. M., Santana, D. L., Dos Santos Oliveira, B. V., Bacci, L. & Albano Araújo, A. P. 2019. Production of extrafloral nectar in the Neotropical shrub *Turnera subulata* mediated by biotic and abiotic factors. Flora 260:151483. Urban & Fischer.

Csardi, G. & Nepusz, T. 2006. The igraph software package for complex network research. InterJournal, Complex Systems 1695.

Del-Claro, K., Rico-Gray, V., Torezan-Silingardi, H. M., Alves-Silva, E., Fagundes, R., Lange, D., Dáttilo, W., Vilela, A. A., Aguirre, A. & Rodriguez-Morales, D. 2016. Loss and gains in ant–plant interactions mediated by extrafloral nectar: fidelity, cheats, and lies. Insectes Sociaux 63:207–221.

Delmas, E., Besson, M., Brice, M.-H., Burkle, L. A., Dalla Riva, G. V., Fortin, M.-J., Gravel, D., Guimarães, P. R., Hembry, D. H., Newman, E. A., Olesen, J. M., Pires, M. M., Yeakel, J. D. & Poisot, T. 2019. Analysing ecological networks of species interactions. Biological Reviews 94:16–39.

Díaz-Castelazo, C., Sánchez-Galván, I. R., Guimarães, P. R., Raimundo, R. L. G. & Rico-Gray, V. 2013. Long-term temporal variation in the organization of an ant-plant network. Annals of botany 111:1285–93.

Domingos, S. S. & Alves Silva, E. 2023. Effect of ants on herbivory levels of *Inga laurina*_: the interplay between space and time in an urban area. Journal of Tropical Ecology 39:e16. Cambridge University Press.

Faber-Langendoen, D., Navarro, G., Willner, W., Keith, D. A., Liu, C., Guo, K. & Meidinger, D. 2020. Perspectives on terrestrial biomes the international vegetation classification. Pp. 1–15 in Goldstein, M. I. & DellaSala, D. A. (eds.). Encyclopedia of the World’s Biomes (1st edition). Elsevier.

Fagundes, R., Dáttilo, W., Ribeiro, S. P., Rico-Gray, V., Jordano, P. & Del-Claro, K. 2017. Differences among ant species in plant protection are related to production of extrafloral nectar and degree of leaf herbivory. Biological Journal of the Linnean Society 122:71–83. Oxford University Press.

Farine, D. R. & Whitehead, H. 2015. Constructing, conducting and interpreting animal social network analysis. Journal of Animal Ecology 84:1144–1163. Wiley/Blackwell (10.1111).

Franco, M. S. & Cogni, R. 2013. Common-garden experiments reveal geographical variation in the interaction among *Crotalaria pallida* (Leguminosae: Papilionideae), *Utetheisa ornatrix* L. (Lepidoptera: Arctiidae), and extrafloral nectary visiting ants. Neotropical Entomology 42:223–229.

González, A. M., Dalsgaard, B. & Olesen, J. M. 2010. Centrality measures and the importance of generalist species in pollination networks. Ecological Complexity 7:36–43.

Guimarães, P. R. 2020. The structure of ecological networks across levels of organization. *Annual Review of Ecology*, Evolution, and Systematics 51:433–460.

Hoffmann, W. A., Geiger, E. L., Gotsch, S. G., Rossatto, D. R., Silva, L. C. R., Lau, O. L., Haridasan, M. & Franco, A. C. 2012. Ecological thresholds at the savanna-forest boundary: how plant traits, resources and fire govern the distribution of tropical biomes. Ecology Letters 15:759–768. John Wiley & Sons, Ltd.

Janzen, D. H. 1969. Allelopathy by myrmecophytes: the ant *Azteca* as an allelopathic agent of Cecropia. Ecology 50:147–153. Wiley Online Library.

Jordán, F., Benedek, Z. & Podani, J. 2007. Quantifying positional importance in food webs: A comparison of centrality indices. Ecological Modelling 205:270–275.

Juárez-Juárez, B., Dáttilo, W. & Moreno, C. E. 2023. Synthesis and perspectives on the study of ant-plant interaction networks: A global overview. Ecological Entomology 48:269–283.

Kersch, M. F. & Fonseca, C. R. 2005. Abiotic factors and the conditional outcome of an ant-plant mutualism. Ecology 86:2117–2126.

Koch, E. B. A., Camarota, F. & Vasconcelos, H. L. 2016. Plant ontogeny as a conditionality factor in the protective effect of ants on a neotropical tree. Biotropica 48:198–205.

Lange, D., Dáttilo, W. & Del-Claro, K. 2013. Influence of extrafloral nectary phenology on ant-plant mutualistic networks in a neotropical savanna. Ecological Entomology 38:463–469.

Leal, L. C., Andersen, A. N. & Leal, I. R. 2015. Disturbance winners or losers? Plants bearing extrafloral nectaries in Brazilian Caatinga. Biotropica 47:468–474. Blackwell Publishing Ltd.

Marazzi, B., Bronstein, J. L. & Koptur, S. 2013. The diversity, ecology and evolution of extrafloral nectaries: current perspectives and future challenges. Ann Bot-London 111:1243–1250.

Marazzi, B., Gonzalez, A. M., Delgado-Salinas, A., Luckow, M. A., Ringelberg, J. J. & Hughes, C. E. 2019. Extrafloral nectaries in Leguminosae: Phylogenetic distribution, morphological diversity and evolution. Australian Systematic Botany 32:409–458. CSIRO PUBLISHING.

Mendes-Silva, I., Queiroga, D., Calixto, E. S., Torezan-Silingardi, H. M. & Del-Claro, K. 2024. Ineffectiveness of ants in protecting two sympatric myrmecophilous plants against endophytic beetles. Austral Ecology 49:1–13. John Wiley & Sons, Ltd.

Miranda, V. S., Rodrigues, L. G., Dutra, S. C., Sobrinho, T. G. & Alves-Araújo, A. 2022. Extrafloral nectaries of an Atlantic Forest conservation area in Southeastern Brazil. Acta Botanica Brasilica 36.

Monique, K., De Souza, G. R., Calixto, E. S. & Silva, E. A. 2022. Temporal variation in the effect of ants on the fitness of myrmecophilic plants: seasonal effect surpasses periodic benefits. Science of Nature 109:1–9.

Morellato, L. P. C. & Oliveira, P. S. 1994. Extrafloral nectaries in the tropical tree *Guarea macrophylla* (Meliaceae). Canadian Journal of Botany 72:157–160.

Moura, R. F., Couto, C. M. V. & Del-Claro, K. 2022. Ant nest distribution and richness have opposite effects on a Neotropical plant with extrafloral nectaries. Ecological Entomology 47:626–635.

Moura, R. F. & Del-Claro, K. 2023. Plants with extrafloral nectaries share indirect defenses and shape the local arboreal ant community. Oecologia 201:73–82. Springer Berlin Heidelberg.

Myers, N., Mittermeier, R. A., Mittermeier, C. G., Da Fonseca, G. A. B. & Kent, J. 2000. Biodiversity hotspots for conservation priorities. Nature 403:853–858.

Navarro, G., Luebert, F. & Molina, J. A. 2023. South American terrestrial biomes as geocomplexes: A geobotanical landscape approach. Vegetation Classification and Survey 4:75–114.

Neves, F. S., Braga, R. F., Araújo, L. S., Campos, R. I. & Fagundes, M. 2012. Differential effects of land use on ant and herbivore insect communities associated with *Caryocar brasiliense* (Caryocaraceae). Revista de Biologia Tropical 60:1065–1073.

Nogueira, A., Guimarães, E., Machado, S. R. & Lohmann, L. G. 2012. Do extrafloral nectaries present a defensive role against herbivores in two species of the family Bignoniaceae in a Neotropical savannas? Plant Ecology 213:289–301.

Nogueira, A., Rey, P. J., Alcántara, J. M., Feitosa, R. M. & Lohmann, L. G. 2015. Geographic mosaic of plant evolution: Extrafloral nectary variation mediated by ant and herbivore assemblages. PLoS ONE 10:1–24.

Oliveira-Filho, L. A., Calixto, E. S., Santos, D. F. B. & Del-Claro, K. 2022. Negative cascading effects of a predatory fly larva on an ant–plant protective mutualism. Arthropod-Plant Interactions 16:373–385. Springer Netherlands.

Oliveira, F. M. P., Câmara, T., Durval, J. I. F., Oliveira, C. L. S., Arnan, X., Andersen, A. N., Ribeiro, E. M. S. & Leal, I. R. 2021. Plant protection services mediated by extrafloral nectaries decline with aridity but are not influenced by chronic anthropogenic disturbance in Brazilian Caatinga. Journal of Ecology 109:260–272.

Oliveira, P. S. 1997. The ecological function of extrafloral nectaries: herbivore deterrence by visiting ants and reproductive output in *Caryocar brasiliense* (Caryocaraceae). Functional Ecology 11:323–330.

Oliveira, P. S. & Leitao-Filho, H. F. 1987. Extrafloral nectaries: their taxonomic distribution and abundance in the woody flora of cerrado vegetation in southeast Brazil. Biotropica 19:140–148.

Oliveira, P. S., Silva, A. F. & Martins, A. B. 1987. Ant foraging on extrafloral nectaries of *Qualea granditlora* (Vochysiaceae) in cerrado: ants as potential antiherbivore agents vegetation. Oecologia 74:228–230.

Oliveira, U., Paglia, A. P., Brescovit, A. D., De Carvalho, C. J. B., Silva, D. P., Rezende, D. T., Leite, F. S. F., Batista, J. A. N., Barbosa, J. P. P. P., Stehmann, J. R., Ascher, J. S., De Vasconcelos, M. F., De Marco, P., Löwenberg-Neto, P., Dias, P. G., Ferro, V. G. & Santos, A. J. 2016. The strong influence of collection bias on biodiversity knowledge shortfalls of Brazilian terrestrial biodiversity. Diversity and Distributions 22:1232–1244.

Oliveira, U., Soares-Filho, B. S., Paglia, A. P., Brescovit, A. D., De Carvalho, C. J. B., Silva, D. P., Rezende, D. T., Leite, F. S. F., Batista, J. A. N., Barbosa, J. P. P. P., Stehmann, J. R., Ascher, J. S., De Vasconcelos, M. F., De Marco, P., Löwenberg-Neto, P., Ferro, V. G. & Santos, A. J. 2017. Biodiversity conservation gaps in the Brazilian protected areas. Scientific Reports 7:9141.

Pebesma, E. & Bivand, R. 2023. Spatial data science: with applications in R. Chapman and Hall/CRC.

Pennington, P. T., Cronk, Q. C. B., Richardson, J. A., Pennington, R. T., Cronk, Q. C. B. & Richardson, J. A. 2004. Introduction and synthesis: Plant phylogeny and the origin of major biomes. Philosophical Transactions of the Royal Society of London. Series B: Biological Sciences 359:1455–1464. Royal Society.

Pereira, R. & Gonçalves, C. 2022. geobr: download official spatial data sets of Brazil_. R package version 1.7.0.

Queiroga, D. Da S. & Moura, R. F. 2017. Positive relation between abundance of pericarpial nectaries and ant richness in *Tocoyena formosa* (Rubiaceae). Sociobiology 64:423–429.

Reu, B., Proulx, R., Bohn, K., Dyke, J. G., Kleidon, A., Pavlick, R. & Schmidtlein, S. 2011. The role of climate and plant functional trade-offs in shaping global biome and biodiversity patterns. Global Ecology and Biogeography 20:570–581. John Wiley & Sons, Ltd.

Roesch, L. F. W., Vieira, F. C. B., Pereira, V. A., Schünemann, A. L., Teixeira, I. F., Senna, A. J. T. & Stefenon, V. M. 2009. The Brazilian Pampa: A fragile biome. Diversity 1:182–198.

Sazima, C., Guimarães, P. R., Dos Reis, S. F. & Sazima, I. 2010. What makes a species central in a cleaning mutualism network? Oikos 119:1319–1325. John Wiley & Sons, Ltd.

Sendoya, S. F., Freitas, A. V. L. & Oliveira, P. S. 2009. Egg-laying butterflies distinguish predaceous ants by sight. The American naturalist 174:134–40. University of Chicago Press.

Shenoy, M., Radhika, V., Satish, S. & Borges, R. 2012. Composition of extrafloral nectar influences interactions between the myrmecophyte *Humboldtia brunonis* and its ant associates. Journal of Chemical Ecology 38:88–99. Springer Netherlands.

Silva, E. A., Anjos, D., Bächtold, A., Lange, D., Maruyama, P. K., Del-Claro, K. & Mody, K. 2020. To what extent is clearcutting vegetation detrimental to the interactions between ants and Bignoniaceae in a Brazilian savanna? Journal of Insect Conservation 24:103–114. Springer International Publishing.

Silva, J. M. C., Leal, I. R. & Tabarelli, M. 2017. Caatinga: The largest tropical dry forest region in South America. Springer. 487 pp.

Simon, M. F., Grether, R., De Queiroz, L. P., Skema, C., Pennington, R. T. & Hughes, C. E. 2009. Recent assembly of the Cerrado, a neotropical plant diversity hotspot, by in situ evolution of adaptations to fire. Proceedings of the National Academy of Sciences 106:20359–20364.

Souza, L. S., Calixto, E. S., Domingos, S. S., Bächtold, A. & Alves-Silva, E. 2024. Ant protection effectiveness in myrmecophytes and extrafloral nectary plants. Journal of Zoology:in press.

Trøjelsgaard, K. & Olesen, J. M. 2016. Ecological networks in motion: micro- and macroscopic variability across scales. Functional Ecology 30:1926–1935.

Weber, M. G. & Keeler, K. H. 2013. The phylogenetic distribution of extrafloral nectaries in plants. Annals of botany 111:1251–61.

Weber, M. G., Porturas, L. D. & Keeler, K. H. 2015. World list of plants with extrafloral nectaries. www.extrafloralnectaries.org.

Wickham, H. 2016. _ggplot2: Elegant graphics for data analysis. Springer-Verlag New York.

Woodward, F. I., Lomas, M. R. & Kelly, C. K. 2004. Global climate and the distribution of plant biomes. Philosophical Transactions of the Royal Society B: Biological Sciences 359:1465. The Royal Society.

Zappi, D. C., Filardi, F. L. R., Leitman, P., Souza, V. C., Walter, B. M. T., Pirani, J. R., Morim, M. P., Queiroz, L. P., Cavalcanti, T. B. & Mansano, V. F. 2015. Growing knowledge: An overview of seed plant diversity in Brazil. Rodriguésia 66:1085–1113. SciELO Brasil.

