## Supplementary Files for "Fabaceae and Cerrado savanna: Two descriptors of Brazilian extrafloral nectary plants" (from another version of this preprint: version 1, 2024-04-09)

Alexandra Bächtold ²

Kleber Del-Claro ^3^

Estevao Alves da Silva ^1^

^1^ Instituto Federal de Educação, Ciência e Tecnologia Goiano, campus Urutaí, Goiás, Brasil

² Independent Researcher. Academic post-doc degree by Universidade Federal de Uberlândia, Uberlândia, MG, Brasil

^3^ Universidade Federal de Uberlândia, Minas Gerais, Brasil

**Supplementary File 1**

**Supplementary File 1**. Box showing the main characteristics of all the six terrestrial Brazilian biomes.

| **Biomes (area and % of territory)** | **Brief features*** |
| --- | --- |
| Amazon  4.196.943 km²  (49.29%) | The largest biome in Brazil and half of the world’s rainforest; home of one third of the Earth’s species; one of the most important carbon sinks in the planet; rains can reach up to 3000 mm per year; phytophysiognomies include mangrove, terra firme, varzea and igapó; low nutrient soils with high content of clay and sand; rains are not evenly distributed in the biome and floods and drought are noted. |
| Caatinga  844,453 km²  (9.92%) | A biome found only in Brazil; Xeric shrubland and thorn forest with vegetation adapted to arid and semi-arid conditions; phytophysiognomies: grassland, xerophytic thorn shrub savanna, and shrub woodland; precipitation up to 1000 mm per year; anomalous climate with high interannual variability in rainfall and droughts; up to 20% of endemic plants. |
| Cerrado  2,036,448 km²  (23.92%) | Hotspot; the riches savanna in the world; dry and wet seasonal variations;  threats include land conversion into agriculture; precipitation up to 1500 mm per year; acidic latosoils soils with high levels of aluminum; part of the vegetation is adapted to fire by possessing meristems below soil and thick barks; phytophysiognomies include grasslands, savanna, cerradão, rocky grasslands and veredas, a moist land. The *Caryocar brasiliense* (Caryocaraceae) (pequi) is recognized by law as a symbol of Cerrado. |
| Mata Atlântica  1,110,182 km²  (13.04%) | Hotspot; Tropical moist forest located along the eastern coast and extending up to 3000 km from south towards the north; presents several climatic conditions and physiognomies such as semi-deciduous, evergreen, deciduous, sub-montane, montane forests; the most threated biome in Brazil as less than 7% of its original cover remains; up to eight thousand endemic plant species; precipitation up to 2000 mm per year; wide range of soils, from eutrophic to dystrophic. |
| Pampa  176,496 km²  (2.07%) | Located at the very southern Brazil; subtropical and temperate climates with four well-characterized seasons; grasslands, shrubs and scarce trees; many plants are xerophytes; main families are Poaceae, Asteraceae, Cyperacea, Fabaceae; fragile sandy soils subjected to water and wind erosion. Threats: livestock production. |
| Pantanal  150,355 km²  (1.76%) | The smallest biome in Brazil; a plateau wetland biome located in a geographic depression and characterized by seasonal cycles of drought and floods; vertebrates are common in the biome, but in general, endemic species are rare; biodiversity threats are caused by fire, pasture, crops, overhunting, invasive species and even unsustainable tourism |

*(Alho, 2011; Eisemberg & Reynolds, 2017; Goldstein & DellaSala, 2020; Junk et al., 2006; Ledru et al., 2016; Marques & Grelle, 2021; Oliveira & Marquis, 2002; Pott et al., 2011; Roesch et al., 2009; Silva & Jr., 2020; Silva et al., 2017).

**References**

Alho, C. J. R. (2011). Biodiversity of the Pantanal: its magnitude, human occupation, environmental threats and challenges for conservation. *Brazilian Journal of Biology*, *71*(1 SUPPL.), 229–232. https://doi.org/10.1590/s1519-69842011000200001

Eisemberg, C. C., & Reynolds, S. J. (2017). *An introduction to wildlife conservation in the Brazilian Amazon: a view from northern Australia*. Brazilian Amazon Field Intensive, Charles Darwin University. https://www.researchgate.net/profile/Andre-Ghidini/publication/329276831_The_Amazon_Environment/links/60392c52a6fdcc37a855891d/The-Amazon-Environment.pdf#page=11

Goldstein, M. I., & DellaSala, D. A. (2020). *Encyclopedia of the world’s biomes*. Elsevier. https://www.sciencedirect.com/referencework/9780128160978/encyclopedia-of-the-worlds-biomes#book-info

Junk, W. J., Da Cunha, C. N., Wantzen, K. M., Petermann, P., Strüssmann, C., Marques, M. I., & Adis, J. (2006). Biodiversity and its conservation in the Pantanal of Mato Grosso, Brazil. *Aquatic Sciences*, *68*(3), 278–309. https://doi.org/10.1007/S00027-006-0851-4/METRICS

Ledru, M.-P., Montade, V., Blanchard, G., & Hély, C. (2016). Long-term spatial changes in the distribution of the Brazilian Atlantic Forest. *Biotropica*, *48*(2), 159–169. https://doi.org/https://doi.org/10.1111/btp.12266

Marques, M. C. M., & Grelle, C. E. V. (2021). *The Atlantic forest: history, biodiversity, threats and opportunities of the mega-diverse forest*. Springer. https://doi.org/https://doi.org/10.1007/978-3-030-55322-7

Oliveira, P. S., & Marquis, R. J. (2002). The cerrados of Brazil: ecology and natural history of a neotropical savanna. In *The cerrados …*. New York: Columbia University Press. https://doi.org/10.1663/0013-0001(2003)057[0656:DFABRE]2.0.CO;2

Pott, A., Oliveira, A. K. M., Damasceno-Junior, G. A., & Silva, J. S. V. (2011). Plant diversity of the Pantanal wetland. *Brazilian Journal of Biology*, *71*(1 SUPPL.), 265–273. https://doi.org/10.1590/s1519-69842011000200005

Roesch, L. F. W., Vieira, F. C. B., Pereira, V. A., Schünemann, A. L., Teixeira, I. F., Senna, A. J. T., & Stefenon, V. M. (2009). The Brazilian Pampa: a fragile biome. *Diversity*, *1*(2), 182–198. https://doi.org/10.3390/d1020182

Silva, J. M. C. da, & Jr., T. E. L. (2020). Caatinga - South America. In M. I. Goldstein & D. A. DellaSala (Eds.), *Encyclopedia of the World’s Biomes* (pp. 554–561). Elsevier. https://doi.org/https://doi.org/10.1016/B978-0-12-409548-9.11984-0

Silva, J. M. C., Leal, I. R., & Tabarelli, M. (2017). *Caatinga: the largest tropical dry forest region in South America*. Springer. https://doi.org/https://doi.org/10.1007/978-3-319-68339-3

**Supplementary File 2**

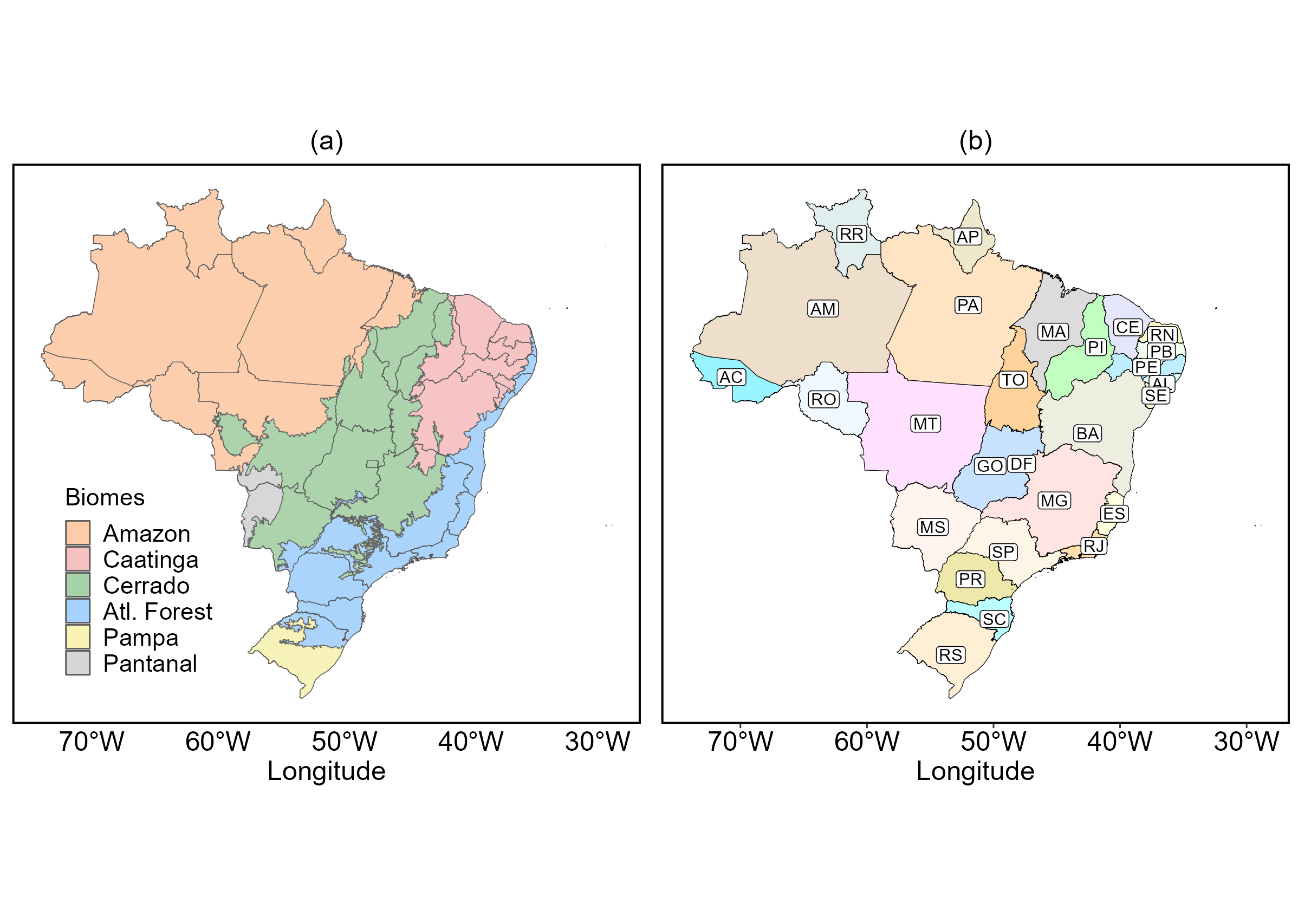

**Supplementary File 2** – Brazilian biomes overlaid with the federated units (26 states plus the capital, the Federal District); (b) Brazilian map showing the geographical frontiers of each federated unit and their abbreviations. Acre (AC), Alagoas (AL)Amapá (AP), Amazonas (AM), Bahia (BA),Ceará (CE), Distrito Federal (DF), Espírito Santo (ES), Goiás (GO), Maranhão (MA), Mato Grosso (MT), Mato Grosso do Sul (MS), Minas Gerais (MG), Pará (PA), Paraíba (PB), Paraná (PR), Pernambuco (PE), Piauí (PI), Rio de Janeiro (RJ), Rio Grande do Norte (RN), Rio Grande do Sul (RS), Rondônia (RO), Roraima (RR), Santa Catarina (SC), São Paulo (SP), Sergipe (SE), Tocantins (TO).

**Supplementary File 3**

**Supplementary File 3.** Families and species of extrafloral nectary plants reportedly visited by ants in Brazilian biomes.

| **Family** | **Species** |
| --- | --- |
| Anacardiaceae | *Anacardium occidentale* |
| Apocynaceae | *Aspidosperma parvifolium* |
| Apocynaceae | *Aspidosperma polyneuron* |
| Apocynaceae | *Aspidosperma pyrifolium* |
| Apocynaceae | *Forsteronia glabrescens* |
| Araceae | *Philodendron cipoense* |
| Araceae | *Philodendron megalophillum* |
| Araceae | *Philodendron myrmecophilum* |
| Asteraceae | *Mikania neurocaula* |
| Asteraceae | *Symphyopappus reticulatus* |
| Bignoniaceae | *Adenocalymma adenophorum* |
| Bignoniaceae | *Adenocalymma albiflorum* |
| Bignoniaceae | *Adenocalymma flaviflorum* |
| Bignoniaceae | *Adenocalymma longilineum* |
| Bignoniaceae | *Adenocalymma moringifolium* |
| Bignoniaceae | *Adenocalymma nodosum* |
| Bignoniaceae | *Adenocalymma peregrinum* |
| Bignoniaceae | *Adenocalymma tanaeciicarpum* |
| Bignoniaceae | *Adenocalymma validum* |
| Bignoniaceae | *Amphilophium elongatum* |
| Bignoniaceae | *Amphilophium parkeri* |
| Bignoniaceae | *Anemopaegma album* |
| Bignoniaceae | *Anemopaegma arvense* |
| Bignoniaceae | *Anemopaegma robustum* |
| Bignoniaceae | *Anemopaegma scabriusculum* |
| Bignoniaceae | *Bignonia aequinoctialis* |
| Bignoniaceae | *Bignonia prieurei* |
| Bignoniaceae | *Cuspidaria sceptrum* |
| Bignoniaceae | *Cybistax antisyphilitica* |
| Bignoniaceae | *Fridericia cf trailli* |
| Bignoniaceae | *Fridericia platyphylla* |
| Bignoniaceae | *Fridericia prancei* |
| Bignoniaceae | *Fridericia triplinervia* |
| Bignoniaceae | *Handroanthus chrysotrichus* |
| Bignoniaceae | *Handroanthus heptaphyllus* |
| Bignoniaceae | *Handroanthus ochraceus* |
| Bignoniaceae | *Jacaranda ulei* |
| Bignoniaceae | *Pachyptera aromatica* |
| Bignoniaceae | *Pleonotoma melioides* |
| Bignoniaceae | *Tabebuia aurea* |
| Bignoniaceae | *Tynanthus panurensis* |
| Bignoniaceae | *Zeyheria tuberculosa* |
| Bixaceae | *Cochlospermum regium* |
| Boraginaceae | *Varronia curassavica* |
| Bromeliaceae | *Aechmea lindenii* |
| Bromeliaceae | *Aechmea nudicaulis* |
| Burseraceae | *Protium heptaphyllum* |
| Burseraceae | *Protium pilosum* |
| Cactaceae | *Pilosocereus tuberculatus* |
| Cactaceae | *Tacinga palmadora* |
| Calophyllaceae | *Caraipa grandifolia* |
| Cannabaceae | *Trema micrantha* |
| Capparaceae | *Cynophalla flexuosa* |
| Caryocaraceae | *Caryocar brasiliense* |
| Chrysobalanaceae | *Hirtella racemosa* |
| Chrysobalanaceae | *Licania humilis* |
| Chrysobalanaceae | *Licania octandra* |
| Combretaceae | *Laguncularia racemosa* |
| Combretaceae | *Terminalia fagifolia* |
| Convovulaceae | *Ipomoea carnea* |
| Convovulaceae | *Ipomoea philomega* |
| Convovulaceae | *Ipomoea regnellii* |
| Costaceae | *Costus scaber* |
| Dennstaedtiaceae | *Pteridium esculentum* |
| Ebenaceae | *Diospyros hispida* |
| Euphorbiaceae | *Aparistimium cordatum* |
| Euphorbiaceae | *Bernardia similis* |
| Euphorbiaceae | *Cnidoscolus bahianus* |
| Euphorbiaceae | *Cnidoscolus quercifolius* |
| Euphorbiaceae | *Croton adamantinus* |
| Euphorbiaceae | *Croton heliotropiifolius* |
| Euphorbiaceae | *Croton jacobinensis* |
| Euphorbiaceae | *Croton nepetifolius* |
| Euphorbiaceae | *Croton tricolor* |
| Euphorbiaceae | *Mabea occidentalis* |
| Euphorbiaceae | *Manihot caerulescens* |
| Euphorbiaceae | *Maprounea brasiliensis* |
| Euphorbiaceae | *Maprounea guianensis* |
| Euphorbiaceae | *Omphalea diandra* |
| Euphorbiaceae | *Sapium glandulosum* |
| Fabaceae | *Albizia pedicellari* |
| Fabaceae | *Anadenanthera colubrina* |
| Fabaceae | *Anadenanthera peregrina* |
| Fabaceae | *Ancistrotropis firmula* |
| Fabaceae | *Bauhinia cheilantha* |
| Fabaceae | *Bauhinia rufa* |
| Fabaceae | *Bionia coriacea* |
| Fabaceae | *Cenostigma microphyllum* |
| Fabaceae | *Cenostigma pyramidale* |
| Fabaceae | *Chamaecrista apoucouita* |
| Fabaceae | *Chamaecrista debilis* |
| Fabaceae | *Chamaecrista desvauxii* |
| Fabaceae | *Chamaecrista ensiformes* |
| Fabaceae | *Chamaecrista mucronata* |
| Fabaceae | *Chamaecrista papillata* |
| Fabaceae | *Chamaecrista ramosa* |
| Fabaceae | *Chamaecrista repens* |
| Fabaceae | *Chloroleucon foliolosum* |
| Fabaceae | *Copaifera langsdorffii* |
| Fabaceae | *Crotalaria micans* |
| Fabaceae | *Crotalaria pallida* |
| Fabaceae | *Dimorphandra mollis* |
| Fabaceae | *Dipteryx odorata* |
| Fabaceae | *Enterolobium gummiferum* |
| Fabaceae | *Erythrina speciosa* |
| Fabaceae | *Hymenaea courbaril* |
| Fabaceae | *Hymenaea stigonocarpa* |
| Fabaceae | *Inga acreana* |
| Fabaceae | *Inga alba* |
| Fabaceae | *Inga blanchetiana* |
| Fabaceae | *Inga calantha* |
| Fabaceae | *Inga capitata* |
| Fabaceae | *Inga cayennensis* |
| Fabaceae | *Inga chartacea* |
| Fabaceae | *Inga densiflora* |
| Fabaceae | *Inga edulis* |
| Fabaceae | *Inga heterophylla* |
| Fabaceae | *Inga lateriflora* |
| Fabaceae | *Inga laurina* |
| Fabaceae | *Inga microcoma* |
| Fabaceae | *Inga paraensis* |
| Fabaceae | *Inga punctata* |
| Fabaceae | *Inga sertulifera* |
| Fabaceae | *Inga sessilis* |
| Fabaceae | *Inga stipularis* |
| Fabaceae | *Inga suaveolens* |
| Fabaceae | *Inga tenuistipula* |
| Fabaceae | *Inga vera* |
| Fabaceae | *Inga vulpina* |
| Fabaceae | *Leucaena leucocephala* |
| Fabaceae | *Machaerium myrianthum* |
| Fabaceae | *Mimosa guilandinae* |
| Fabaceae | *Mimosa tenuiflora* |
| Fabaceae | *Parkia pendula* |
| Fabaceae | *Piptadenia retusa* |
| Fabaceae | *Piptadenia uaupensis* |
| Fabaceae | *Pityrocarpa moniliformis* |
| Fabaceae | *Plathymenia reticulata* |
| Fabaceae | *Pterodon pubescens* |
| Fabaceae | *Senegalia bahiensis* |
| Fabaceae | *Senegalia multipinnata* |
| Fabaceae | *Senegalia piauhiensis* |
| Fabaceae | *Senegalia polyphylla* |
| Fabaceae | *Senna cathartica* |
| Fabaceae | *Senna macranthera* |
| Fabaceae | *Senna pendula* |
| Fabaceae | *Senna reniformis* |
| Fabaceae | *Senna rizzinii* |
| Fabaceae | *Senna rugosa* |
| Fabaceae | *Senna splendida* |
| Fabaceae | *Senna velutina* |
| Fabaceae | *Stryphnodendron adstringens* |
| Fabaceae | *Stryphnodendron polyphyllum* |
| Fabaceae | *Stryphnodendron pulcherrimum* |
| Fabaceae | *Stryphnodendron rotundifolium* |
| Fabaceae | *Tachigali venusta* |
| Gentianaceae | *Calolisianthus speciosus* |
| Gentianaceae | *Chelonanthus viridiflorus* |
| Gesneriaceae | *Codonanthe opsisulei* |
| Gesneriaceae | *Codonanthopsis calcarata* |
| Gesneriaceae | *Codonanthopsis crassifolia* |
| Gesneriaceae | *Codonanthopsis uleana* |
| Lamiaceae | *Aegiphila integrifolia* |
| Lamiaceae | *Aegiphila verticillata* |
| Lecythidaceae | *Gustavia augusta* |
| Loganiaceae | *Strychnos panurensis* |
| Lythraceae | *Lafoensia pacari* |
| Malpighiaceae | *Banisteriopsis angustifolia* |
| Malpighiaceae | *Banisteriopsis argyrophylla* |
| Malpighiaceae | *Banisteriopsis campestris* |
| Malpighiaceae | *Banisteriopsis laevifolia* |
| Malpighiaceae | *Banisteriopsis malifolia* |
| Malpighiaceae | *Banisteriopsis muricata* |
| Malpighiaceae | *Banisteriopsis stellaris* |
| Malpighiaceae | *Heteropterys byrsonimifolia* |
| Malpighiaceae | *Heteropterys pteropetala* |
| Malpighiaceae | *Heteropterys umbellata* |
| Malpighiaceae | *Peixotoa parviflora* |
| Malpighiaceae | *Peixotoa tomentosa* |
| Malpighiaceae | *Tetrapterys microphylla* |
| Malvaceae | *Byttneria benensis* |
| Malvaceae | *Eriotheca gracilipes* |
| Malvaceae | *Gossypium hirsutum* |
| Malvaceae | *Talipariti pernambucense* |
| Meliaceae | *Guarea macrophylla* |
| Meliaceae | *Trichilia pallida* |
| Myrsinaceae | *Myrsine guianensis* |
| Myrtaceae | *Psidium australe* |
| Myrtaceae | *Psidium grandifolium* |
| Ochnaceae | *Ouratea castaneifolia* |
| Ochnaceae | *Ouratea hexasperma* |
| Ochnaceae | *Ouratea spectabilis* |
| Orchidaceae | *Epidendrum cinnabarinum* |
| Orchidaceae | *Epidendrum denticulatum* |
| Orchidaceae | *Epidendrum orchidiflorum* |
| Passifloraceae | *Passiflora alata* |
| Passifloraceae | *Passiflora ceratocarpa* |
| Passifloraceae | *Passiflora coccinea* |
| Passifloraceae | *Passiflora edulis* |
| Passifloraceae | *Passiflora quadrangularis* |
| Piperaceae | *Peperomia macrosthachya* |
| Primulaceae | *Myrsine monticola* |
| Rhamnaceae | *Gouania frangulifolia* |
| Rubiaceae | *Cordiera elliptica* |
| Rubiaceae | *Cordiera macrophylla* |
| Rubiaceae | *Cordiera sessilis* |
| Rubiaceae | *Declieuxia fruticosa* |
| Rubiaceae | *Palicourea rigida* |
| Rubiaceae | *Remijia amazonica* |
| Rubiaceae | *Tocoyena formosa* |
| Salicaceae | *Banara guianensis* |
| Sapindaceae | *Serjania clematidea* |
| Simaroubaceae | *Simarouba amara* |
| Smilacaceae | *Smilax polyantha* |
| Solanaceae | *Markea longiflora* |
| Solanaceae | *Solanum stramonifolium* |
| Turneraceae | *Turnera cearensis* |
| Turneraceae | *Turnera subulata* |
| Turneraceae | *Turnera trioniflora* |
| Verbenaceae | *Citharexylum myrianthum* |
| Verbenaceae | *Stachytarpheta glabra* |
| Vochysiaceae | *Qualea grandiflora* |
| Vochysiaceae | *Qualea multiflora* |
| Vochysiaceae | *Qualea parviflora* |

**Supplementary File 4**

**Supplementary File 4.** Frequency in papers and richness of families of extrafloral nectary plants sampled in Brazilian biomes. The sampling covered 170 published papers.

| **Family** | **Richness** | **Papers** |
| --- | --- | --- |
| Fabaceae | 76 | 48 |
| Bignoniaceae | 32 | 12 |
| Euphorbiaceae | 15 | 13 |
| Malpighiaceae | 13 | 36 |
| Rubiaceae | 7 | 15 |
| Passifloraceae | 5 | 13 |
| Malvaceae | 4 | 12 |
| Apocynaceae | 4 | 2 |
| Gesneriaceae | 4 | 2 |
| Vochysiaceae | 3 | 29 |
| Ochnaceae | 3 | 19 |
| Turneraceae | 3 | 8 |
| Convovulaceae | 3 | 6 |
| Chrysobalanaceae | 3 | 4 |
| Araceae | 3 | 3 |
| Orchidaceae | 3 | 3 |
| Burseraceae | 2 | 4 |
| Verbenaceae | 2 | 4 |
| Cactaceae | 2 | 3 |
| Lamiaceae | 2 | 3 |
| Combretaceae | 2 | 2 |
| Gentianaceae | 2 | 2 |
| Meliaceae | 2 | 2 |
| Myrtaceae | 2 | 2 |
| Solanaceae | 2 | 2 |
| Asteraceae | 2 | 1 |
| Bromeliaceae | 2 | 1 |
| Caryocaraceae | 1 | 30 |
| Lythraceae | 1 | 9 |
| Smilacaceae | 1 | 4 |
| Cannabaceae | 1 | 3 |
| Costaceae | 1 | 3 |
| Anacardiaceae | 1 | 2 |
| Boraginaceae | 1 | 2 |
| Capparaceae | 1 | 2 |
| Lecythidaceae | 1 | 2 |
| Loganiaceae | 1 | 2 |
| Myrsinaceae | 1 | 2 |
| Rhamnaceae | 1 | 2 |
| Sapindaceae | 1 | 2 |
| Simaroubaceae | 1 | 2 |
| Bixaceae | 1 | 1 |
| Calophyllaceae | 1 | 1 |
| Dennstaedtiaceae | 1 | 1 |
| Ebenaceae | 1 | 1 |
| Piperaceae | 1 | 1 |
| Primulaceae | 1 | 1 |
| Salicaceae | 1 | 1 |

**Supplementary File 5**

**Supplementary File 5**. Number of species and families of extrafloral nectaried plants in Brazilian biomes. The numbers in parentheses indicate the number of exclusive species and families. Total richness = 224 species; total families = 48. Superscript letters indicate statistically significant differences according to chi-squared tests.

| **Biomes** | **Richness** | **% richness** | **Excl. species** | **Families** | **% families** | **Excl. families** |
| --- | --- | --- | --- | --- | --- | --- |
| Cerrado | 90 ^a^ | 40 | 82 ^a^ | 23 ^a^ | 48 | 12 ^a^ |
| Amazon | 70 ^a^ | 31 | 65 ^a^ | 24 ^a^ | 50 | 9 ^a^ |
| Caatinga | 39 ^b^ | 17 | 34 ^b^ | 11 ^a^ | 23 | 4 ^a^ |
| Atl. Forest | 37 ^b^ | 17 | 29 ^b^ | 17 ^a^ | 35 | 4 ^a^ |
| Pampa | 2 ^c^ | 1 | 1 ^c^ | 1 ^b^ | 2 | 0 ^b^ |

**Supplementary File 6**

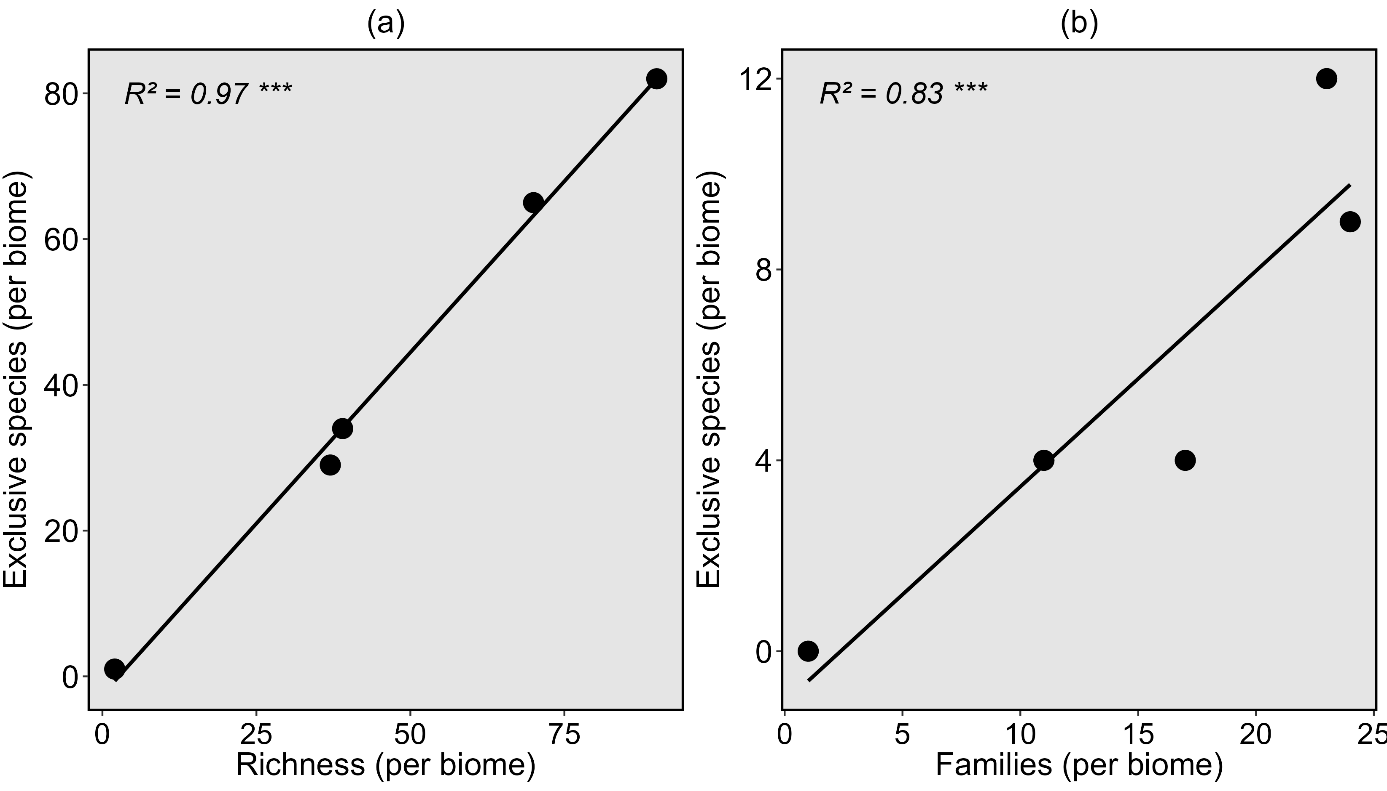

**Supplementary File 6.** Positive relationship between the number of **(a)** exclusive species and the total richness; **(b)** the same rationale, but for families. The sample size in both cases is low, but the slope reveals a clear pattern of data.
